# Murine norovirus capsid plasticity – Glycochenodeoxycholic acid stabilizes P-domain dimers and triggers escape from antibody recognition

**DOI:** 10.1101/2021.02.27.433148

**Authors:** Robert Creutznacher, Thorben Maaß, Jasmin Dülfer, Clara Feldmann, Veronika Hartmann, Jan Knickmann, Leon Torben Westermann, Thomas J. Smith, Charlotte Uetrecht, Alvaro Mallagaray, Thomas Peters, Stefan Taube

## Abstract

The murine norovirus (MNV) capsid protein is the target for various neutralizing antibodies binding to distal tips of its protruding (P)-domain. The bile acid glycochenodeoxycholic acid (GCDCA), an important co-factor for murine norovirus (MNV) infection, has recently been shown to induce conformational changes in surface-loops and a contraction of the virion. Here, we employ protein NMR experiments using stable isotope labeled MNV P-domains to shed light on underlying molecular mechanisms. We observe two separate sets of NMR resonance signals for P-domain monomers and dimers, permitting analysis of the corresponding exchange kinetics. Unlike human norovirus GII.4 P-dimers, which exhibit a half-life in the range of several days, MNV P-dimers are very short lived with a half-life of about 17 s. Addition of GCDCA shifts the equilibrium towards the dimeric form by tightly binding to the P-dimers. In MNV virions GCDCA-mediated stabilization of the dimeric arrangement of P-domains generates a more ordered state, which in turn may entropically assist capsid contraction. Numerous long-range chemical shift perturbations (CSPs) upon addition of GCDCA reflect allosteric conformational changes as a feature accompanying dimer stabilization. In particular, CSPs indicate rearrangement of the E’F’ loop, a target for various neutralizing antibodies. Indeed, treating MNV virions with GCDCA prior to neutralizing antibody exposure abolishes neutralization. These findings advance our understanding of GCDCA-induced structural changes of MNV capsids and experimentally support an intriguing viral immune escape mechanism relying on GCDCA-triggered conformational changes of the P-dimer.

**Significance Statement:** This study sheds light on the role of glycochenodeoxycholic acid (GCDCA) in promoting murine norovirus (MNV) infection and immune escape. Binding of GCDCA to the dimeric P-domain has been well characterized by crystallography and cryo EM studies, showing that upon GCDCA binding, a 90° rotation of the P-domain occurs, which results in its collapse onto the underlying shell of the virus. Our NMR experiments now reveal P-dimer stability as a new dimension of plasticity of MNV capsids and suggest that capsid contraction is entropically assisted. Conformational changes as a feature of P-dimer stabilization eliminate recognition by neutralizing antibodies, no longer being able to prevent infection. These findings highlight key differences between human and MNV capsid structures, promote our understanding of MNV infection on a molecular level, and reveal a novel immune escape mechanism.

## Introduction

Noroviruses are non-enveloped, positive strand RNA viruses, belonging to the family of *Caliciviridae*. Human noroviruses (HuNoVs) are responsible for more than 600 million cases of viral gastroenteritis worldwide annually (1–3). Infections pose a significant burden to health care systems and neither licensed vaccines nor antiviral therapies are currently available. Cell culture systems for HuNoV became only recently available though not trivial to implement (4, 5). Murine noroviruses (MNV), on the other hand, share the enteric tropism with their human counterpart and can be easily studied in cell culture and their native small animal host (6, 7). All noroviruses share a common capsid structure containing 180 copies of the major capsid protein VP1 forming T = 3 icosahedral particles. With the overall capsid structures of HuNoV and MNV being similar one would expect comparable viral strategies to enter host cells. However, significant differences exist particularly during host cell entry. For MNV members of the CD300 receptor family function as a proteinaceous entry receptor (8), while HuNoV require histo blood group antigens (HBGA) for infection. Interestingly both, HuNoV and MNV, bind bile acids, in particular glycochenodeoxycholic acid (GCDCA). Although GCDCA binds to HuNoV and MNV capsids at very different locations and with large differences in binding affinity, GCDCA can promote infection in either species, for HuNoV likely in a strain-specific manner (5, 9–11). Exciting results from cryo-EM studies showed that norovirus capsids exist in at least two major forms, one with the P-domain hovering 10 to 15 Å above the shell domain representing an extended conformation, and another one with the P-domain tightly associated with the shell surface representing a contracted conformation (12–17). For MNV, bile salts were key mediators to drive the capsids into the contracted form, with GCDCA being the most effective (13). However, the mechanism of how the capsid is forced into the contracted form is still elusive.

Crystal structures of MNV P-dimers complexed with the receptor CD300lf in the absence (pdb 6C6Q) and in the presence (pdb 6E47) of GCDCA are available and virtually identical (RMSD 0.495 Å) (10) but the cryo-EM structure of the MNV virion complexed with GCDCA revealed a rotation and contraction of the P domain onto the shell surface compared to the apo structure (13, 17). Interestingly, the crystal structure of the apo form of the MNV-1 P-dimer (pdb 3LQ6) differs from all other available crystal structures complexed either with GCDCA, Fab fragments, or nanobodies in their surface loop structures. In this apo structure of the MNV P-domain, the A’B’ loop (aa 299-301) and the E’F’ loop (aa 379-388) are found in two different orientations, an “open” and a “closed” conformation (18). In the open conformation, the C’D’ loop (aa 342-351) blocks access to the GCDCA binding site at the lateral dimer interface (10) as is illustrated in Fig. 1.

**Fig. 1:**
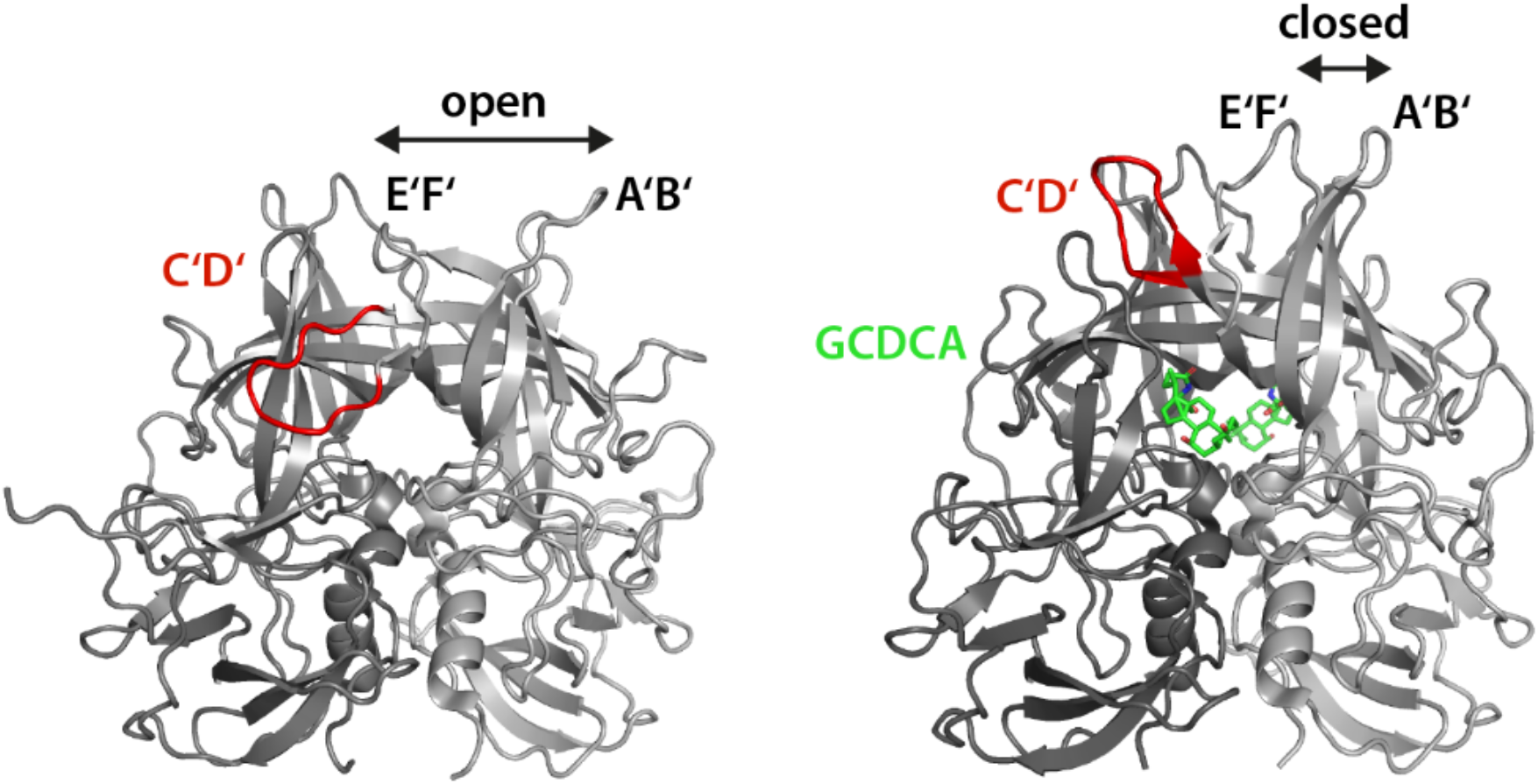
Comparison of structures of the MNV-1 P-dimer in the absence (left, pdb 3LQ6) and presence of GCDCA (right, pdb 6E47). The C’D’ loop can block access to the GCDCA binding pocket in the unbound state of the protein. GCDCA (green) binding displaces the C’D’ loop and triggers concerted movements of the distal A’B’ and E’F’ loops, thereby rearranging the exposed parts of the P-domain into a closed structure.

Antibody escape mutations at the tip of the A’B’ and E’F’ loop have been described for the monoclonal antibodies MAb A62.2 and 2D3 and are thought to be linked to allostery-like mechanisms in the viral capsid (19). Molecular dynamics assisted flexible fitting simulations further suggested that escape mutations, such as V339I stabilize the P-domain dimer interface, thereby affecting conformational features of the P-domain and MAb 2D3 binding. Cryo-EM structures suggested that both MAbs bind to same site of the P domain making contacts to the E’F’ loop, however, none of the natural 2D3 escape mutations (D348E and V339I) were functional for A6.2 and none of the A6.2 escape mutations (A382, D385, V378, L386) were functional for 2D3, suggesting that structural changes rather than point contacts are causing the escape (13, 19). As binding of GCDCA at the dimer interface significantly rearranges the surface loops A’B’, C’D’ and E’F’, compared to the apo structure (pdb 3LQ6), we hypothesized that GCDCA binding impacts MAb neutralization.

In our present study, we focus on the role of bile acids on MNV P-domain plasticity and employed protein NMR experiments in combination with other biophysical techniques, such as native mass spectrometry and analytical size exclusion chromatography (20), to identify modes of flexibility on a molecular level and their impact on immune escape.

## Results

### Synthesis of MNV P-domains and purification

For our studies we used P-domains of three different MNV strains, CW1 (18), CR10 (10), and MNV07 (21), sharing more than 90% sequence identity (Fig. S1 and Table S1). Crystal structure data are available for several P-domains including the strains CW1 and CR10 (10, 18, 22), allowing to correlate NMR data with structural features. This is particularly interesting and informative for data from NMR binding experiments employing stable isotope labeled proteins (23–28). Most experiments in the present study were performed with CW1 P-domains. Compared to prior work (29), we used constructs that were truncated at the C-terminus. Our standard protocols for protein biosynthesis and purification of HuNoV P-domains (30, 31) led to irreversible unfolding and protein aggregation and were not applicable to MNV P-domains (Fig. S2a). MNV P-domains were unstable at neutral pH, and, therefore, purification was performed in acetate buffer at acidic pH. Purification at acidic pH values prevented aggregation and provided properly folded protein (Fig. S2b, cf. Materials and Methods). Analysis of the thermostability of P-domains underlines the importance of adjusting the pH value as the relative stability quickly decreases with increasing pH values above pH 6 (Fig. S2c). The modified protocol was further optimized for the preparation of [*U*-^2^H,^15^N] labeled and of specifically ^13^C-methyl (MIL^proS^V^proS^A) labeled MNV P-domain samples, allowing ^1^H,^15^N TROSY HSQC and Methyl TROSY based chemical shift perturbation experiments, respectively, into P-domain dimerization and ligand binding.

### NMR spectra uncover slow, concentration-dependent monomer-dimer interconversion

Recording a series of ^1^H,^15^N TROSY HSQC spectra of [*U*-^2^H,^15^N] labeled CW1 P-domain samples for protein concentrations ranging from 25 to 200 μM yields spectra with obvious concentration dependent changes. Besides the overall increase in backbone H^N^ cross peak intensities due to improved signal-to-noise ratios at higher concentrations, there are two sets of cross peaks with concentration dependent relative intensities. New cross peaks emerge with increasing P-domain concentrations. Concomitantly, another set of cross peaks decreases in relative intensity as this is exemplified in Figs. 2a and 2b. This concentration dependent behavior is also noticeable in complementary biophysical experiments. Concentration dependent size exclusion chromatography (SEC) runs with P-domain concentrations ranging from 250 nM to 270 μM reflect an increasing molecular weight (Fig. 2c), and native mass spectrometry (MS) experiments directly show that MNV P-domains exist as a mixture of monomers and dimers (Fig. 2d). Normalized SEC curves show increased tailing at higher concentrations indicating monomer-dimer exchange on the time scale of the chromatographic separation (20). Therefore, the two sets of cross peaks in the HSQC spectrum reflect an equilibrium between monomeric and dimeric P-domains. It follows that the peaks increasing in intensity with increasing protein concentration must be assigned to P-domain dimers, and the peaks decreasing in intensity to P-domain monomers.

**Fig. 2:**
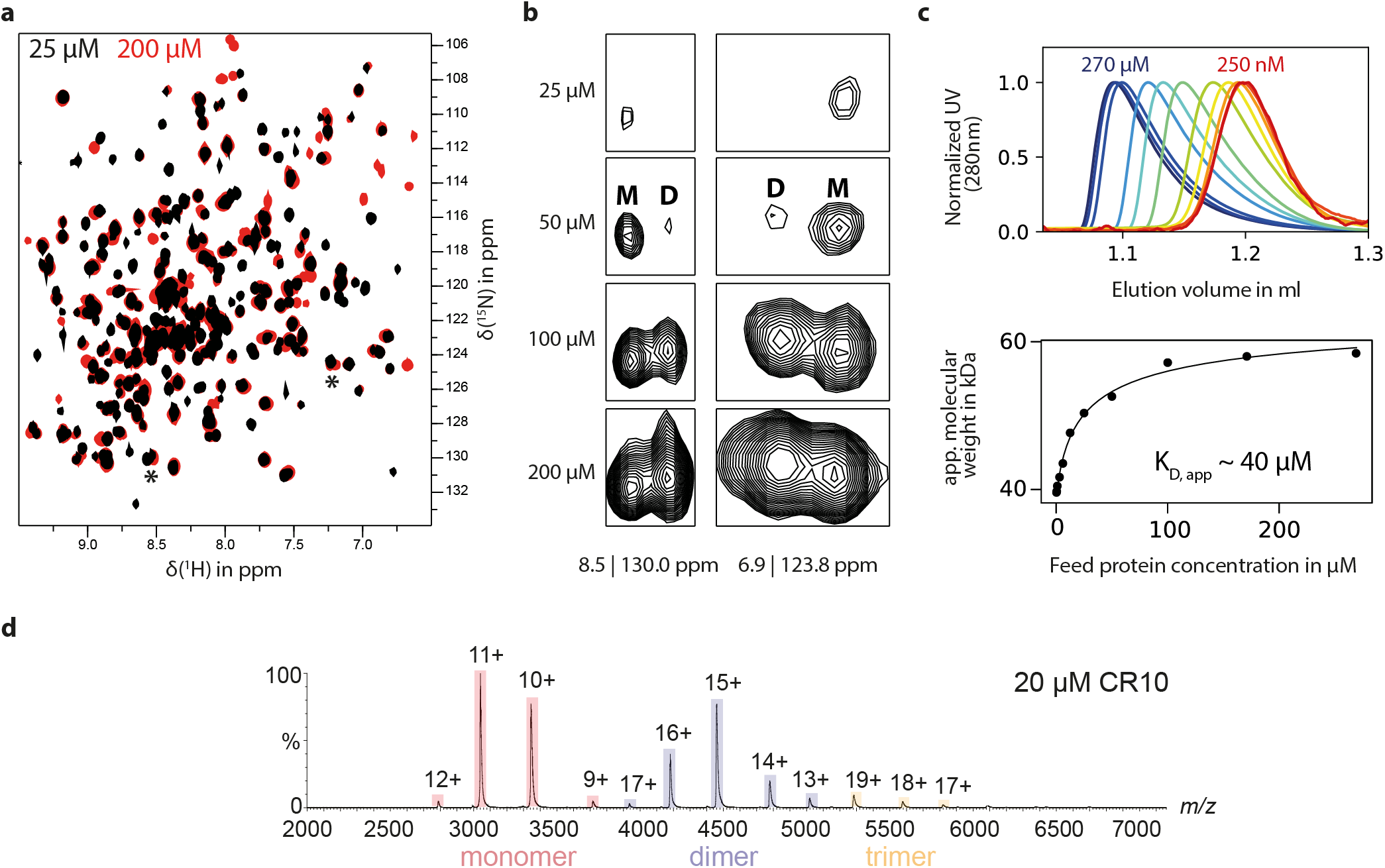
Concentration-dependent dimerization of the MNV P-domain in solution. ^1^H,^15^N TROSY HSQC spectra of [*U*-^2^H,^15^N] labeled MNV CW1 P-domains show protein concentration-dependent changes (**a**, black: 25 μM, red: 200 μM). Several new backbone NH signals appear with increasing protein concentration while others decrease in relative intensity. These signals can be assigned to originate from the dimer (D) or the monomer (M), respectively. Two such signal pairs are marked by asterisks and are shown in **b**. CW1 P-domains have been subjected to size exclusion chromatography at increasing concentrations (**c**), ranging from 250 nM (red) to 270 μM (blue). The elution volume strongly depends on protein concentration and can be associated with an increase of apparent molecular weight from 39.6 kDa (monomer) to 58.4 kDa (dimer). The normalized chromatograms reveal a noticeable peak tailing at higher concentrations, indicating ongoing monomer-dimer exchange on the timescale of the chromatographic separation. Fitting the concentration-dependent apparent molecular weight against the law of mass action yields an apparent *K_D_* value of the dimerization reaction of 40 μM. **(d)** Native MS spectra confirming monomer-dimer equilibrium in a highly similar strain CR10.

Fitting the law of mass action to normalized SEC curves (cf. Eq. 1 in Materials and Methods) and to native MS spectra intensities (cf. Eq. 9 in Materials and Methods) allows estimation of dissociation constants *K_D_* for the P-domain monomer-dimer equilibrium. For CW1 P-dimers an apparent dissociation constant *K_D1_* of 40 μM is obtained from SEC (Fig. 2c), and for CR10 P-dimers a value of 32 μM is obtained from native MS (Fig. S3). Increasing intensities of backbone H^N^ cross peaks were also used to estimate the dissociation constant for CW1 P-dimers, yielding a value of 31 μM. These dissociation constants are considered “apparent” as each experimental approach suffers from certain limitations. In addition to providing estimates of dissociation constants NMR spectra also contain information about the time scale of monomer-dimer exchange. From Fig. 2b it is obvious that monomer and dimer peaks are separate, revealing slow exchange on the NMR chemical shift time scale. Measurement of distances between monomer and dimer cross peaks directly yields an upper limit for the exchange rate constant *k_ex_* of 10 s^−1^, indicating that the “true” value is at least about an order of magnitude smaller. This matches well with the analysis of peak tailing in analytical SEC runs, yielding an estimate for the dissociation rate constant *k_ex_* of 0.03 s^−1^ (cf. Eqs. 2–6 in Materials and Methods).

As an alternative to uniform ^2^H,^15^N backbone labeling we have followed specific ^13^C side chain methyl group labeling strategies (23, 27, 28). Subjecting specifically ^13^C-methyl (MIL^proS^V^proS^A) labeled CW1 P-domain to methyl TROSY experiments allowed to directly monitor the monomer-dimer exchange process. The advantage of ^13^C methyl group labeling is the relative ease with which (partial) assignments of side chain methyl group signals can be obtained, provided high resolution crystal structure data for the protein in question are available. Analysis of a 4D HMQC-NOESY-HMQC spectrum (32, 33) of a MIL^proS^V^proS^A labeled sample of the CW1 P-domain in the presence of saturating amounts of GCDCA in combination with crystal structure data for the protein complexed with GCDCA (10) allowed assignment of about 50% of all ^13^C-methyl resonance signals (Fig. S4). As will be discussed below in detail, chemical shifts of H^N^ backbone protons and of ^13^C methyl groups of the apo form of CW1 P-domains differ notably from the corresponding chemical shifts in the GCDCA-bound form. For CW1 P-domains chemical exchange between apo and bound form is slow on the chemical shift time scale resulting in corresponding peaks at distinct chemical shifts. Under these so called slow-exchange conditions different monomer-dimer ratios caused by changing overall protein concentrations are reflected only by relative intensities of the corresponding cross peaks. There is no continuous change of resonance signal positions as a consequence of averaged chemical shifts as this would be observed under fast exchange conditions (34). Therefore, it is not trivial to transfer the assignment from the CW1 P-domain complexed with GCDCA to the apo form, and, for now, only a subset of signals of the apo form has been assigned. However, it is of course possible to monitor concentration dependent changes of ^13^C methyl group signals of the apo form without having any assignments. The two-dimensional data set, consisting of concentration dependent methyl TROSY spectra, was simulated using direct quantum mechanical simulations and fitting regions of interest to a kinetic model for the exchange process (35). This procedure provides corresponding rate and dissociation constants (cf. Fig. S5). Assuming a two-site exchange model we obtained a dissociation rate constant *k_off_* of 0.035 s^−1^ and a dissociation constant *K_D1_* of 12.5 μM, corresponding to an association rate constant *k_on_* of 2.8 × 10^3^ M^−1^s^−1^ for the CW1 P-domain monomer-dimer exchange process.

### GCDCA promotes P-domain dimerization

When analyzing samples of CW1 P-domains in the presence of saturating amounts of GCDCA with SEC, elution profiles are insensitive to the protein concentration, and the elution volume translates into a molecular weight of ca. 64 kDa as expected for P-domain dimers (Fig. 3a). Simultaneously, the thermostability of CW1 P-domains increases significantly with increasing GCDCA concentrations (Fig. 3b). No such effects are observed in the presence of other non- or weakly binding bile acids such as cholic acid (CA), taurocholic acid (TCA) or taurochenodeoxycholic acid (TCDCA) (Fig. S6). This increased thermostability in the presence of GCDCA is also seen in infectivity assays with MNV virions (Fig. 3c). A higher infectivity is maintained at temperatures above ca. 45°C in the presence of GCDCA. The observed increase of the apparent molecular weight of P-dimers upon GCDCA binding is also reflected by TRACT NMR experiments (36) demonstrating a significant increase of the motional correlation time τ_c_ from 35 to 50 ns, as expected for an increase of molecular weight upon GCDCA binding (Fig. S7). Likewise, measurement of longitudinal (T_1_) and transverse (T_1ρ_, T_2_) relaxation times yielded T_1_/T_2_ ratios reflecting rotational and internal motions on the ps - ns time scale. The average values significantly increase upon addition of GCDCA, indicating an increase in the molecular tumbling time and thus also in molecular weight (Fig. S8). This underscores that P-domain dimerization is significantly promoted by addition of GCDCA.

**Fig. 3.:**
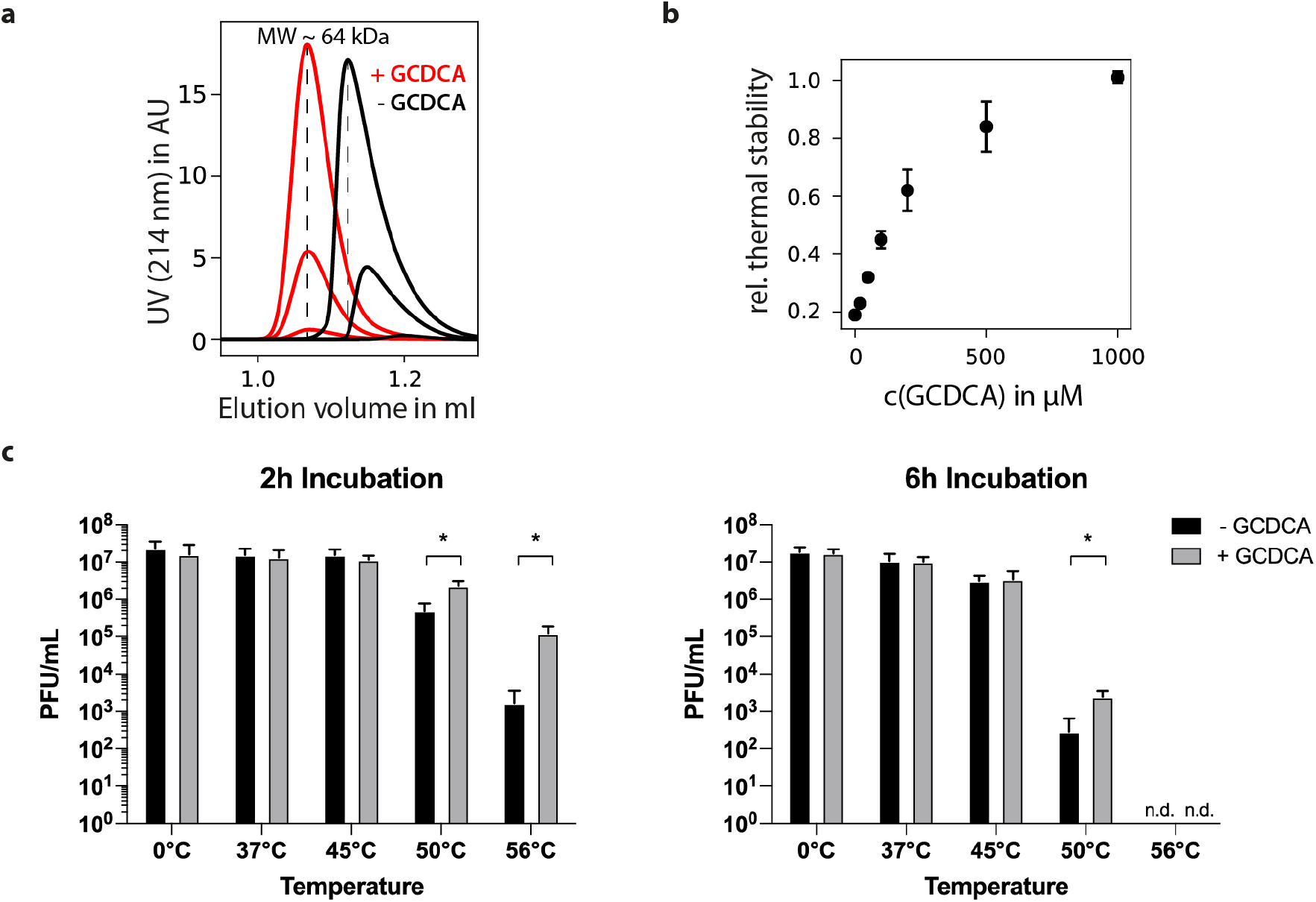
The MNV P-domain readily dimerizes in presence of GCDCA. Size exclusion chromatography of the MNV.CW1 P-domain in presence and absence of GCDCA (**a**, red and black curves, respectively) reveals a major change in elution behavior. The presence of saturating amounts of GCDCA shifts the apparent molecular weight of the P-domain to that of the dimer (64 kDa) regardless of the protein concentration used. Six representative chromatograms are shown with a protein concentration range of 250 nM to 50 μM. The disappearance of the protein concentration-dependent elution behavior indicates a dramatic stabilization of the dimeric protein in presence of its ligand. **(b)** The thermal stability of the CW1 P-domain increases with increasing GCDCA concentrations. Isothermal denaturation for 30 min at 45 °C and analysis of non-denatured P-domain via HIC shows complete protection with 1mM GCDCA when compared to a non-heat-treated control. HIC experiments were performed as duplicates. The respective percentage of deviation is given as error bars. **(c)** Thermal stability profile of MNV-1.CW3 virions in the presence and absence of the bile acid GCDCA. 1 × 10^7^ plaque forming unit (PFU) of MNV-1.CW3 were incubated in the presence or absence of 500 μM GCDCA for 2 or 6 hours at temperatures between 0 and 56 °C. Duplicate plaque assays were performed for three independent assays. Bars represent the mean (n=3) ± standard deviation (SD). Statistical analysis was performed using the unpaired *t*-test (p-value < 0.05). The * indicates significance (*p*<0.05) comparing titers obtained in the presence or absence of GCDCA. (n.d. = no detectable titer).

To gain insight into the molecular basis of GCDCA assisted P-domain dimerization we performed chemical shift perturbation (CSP) NMR experiments. CSPs of backbone H^N^ cross peaks and ^13^C-methyl group signals of amino acid side chains upon addition of GCDCA were followed in ^1^H,^15^N TROSY HSQC spectra of [*U*-^2^H,^15^N] labeled CW1 P-domain and in methyl TROSY spectra of MIL^proS^V^proS^A labeled CW1 P-domain, respectively (Fig. 4). As alluded to above, for the side chain methyl groups a partial assignment is available (Fig. S4) and, therefore, CSPs can be used to localize conformational changes or binding sites. It becomes clear that the presence of saturating amounts of GCDCA leads to dramatic changes of the entire set of signals in the ^1^H,^15^N TROSY HSQC spectrum as well as in the methyl TROSY spectrum (Figs. 4a and 4b). For the methyl TROSY spectrum a significant number of CSPs can be assigned to individual amino acid side chain methyl groups (Figs. 4c and 4d). In addition, peaks originating from P-domain monomers disappear. Signals in the methyl TROSY spectrum can be classified according to the magnitude of the CSPs (CSPs are reported as Euclidian distances Δ*v* (37)). A first set of signals experiences substantial CSPs with Δ*v* much larger than ca. 20 Hz. Due to these large changes, it is difficult to trace such signals back to the apo form. A second set of signals displays CSPs with Δ*v* values in an intermediate range of 10 to 16 Hz, and a third set of signals is classified as weak with Δ*v* values ranging from 6 to 10 Hz (Fig. 4c and 4d). CSPs below a threshold of 6 Hz were not considered here. This classification of CSPs is arbitrary but follows published recommendations (37). Not surprisingly, methyl side chains of amino acids close to the GCDCA binding pocket are most affected (Fig. 4c). Inspection of ^1^H,^15^N TROSY HSQC spectra of P-domain titrated with GCDCA yields a matching data set with some signals in slow and some in fast chemical exchange on the chemical shift time scale (Fig. S9). This allows to estimate the exchange rate constant *k_ex_* for binding of GCDCA to MNV P-dimers placing it in a range between 30 and 60 s^−1^.

**Fig. 4.:**
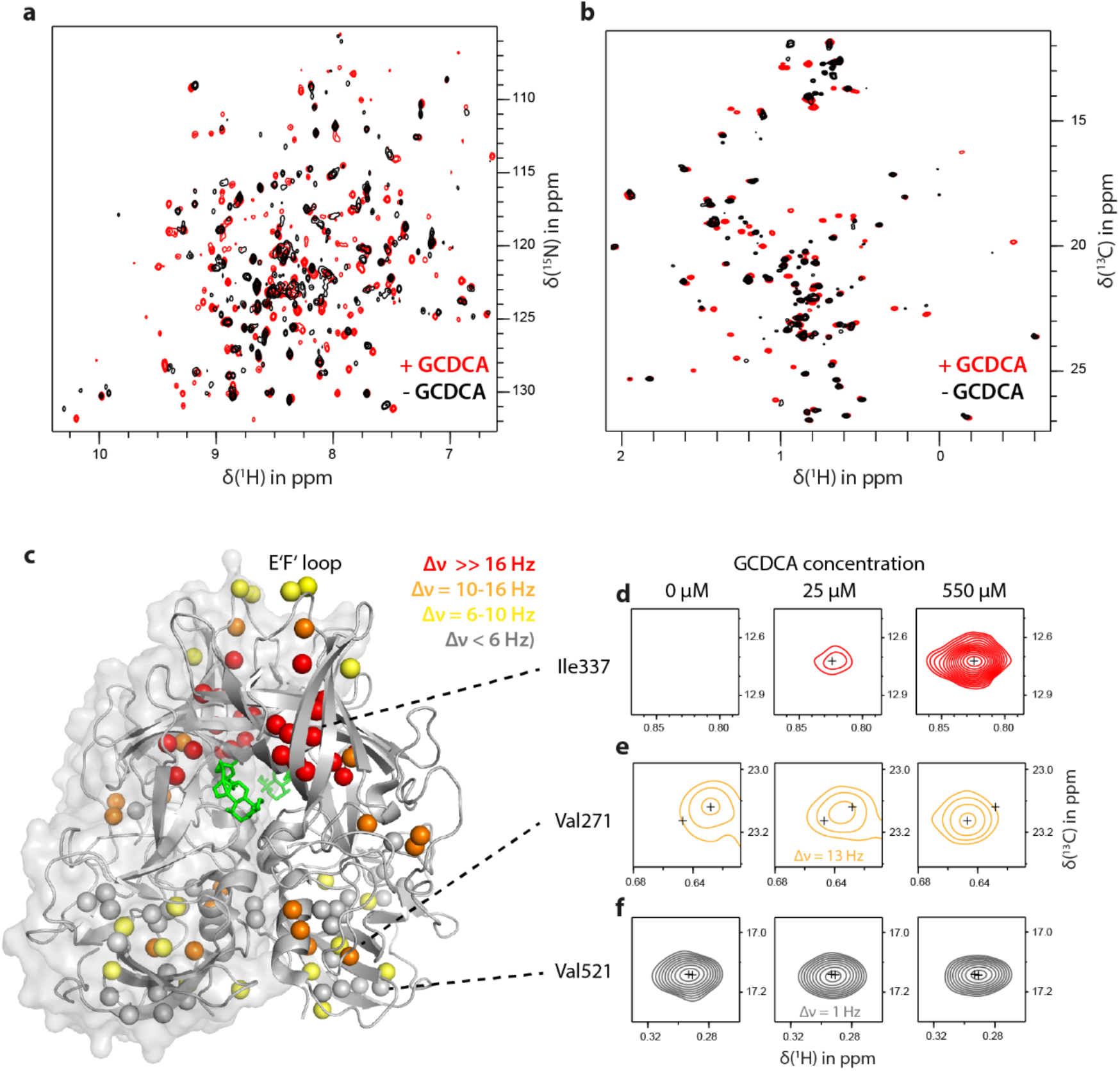
Chemical shift perturbations in MNV CW1 P-domains upon GCDCA binding. ^1^H,^15^N TROSY HSQC and methyl TROSY spectra of 100 μM backbone labeled (**a**) or 50 μM methyl group-labeled (**b**) MNV P-domains titrated with GCDCA (black and red spectra, respectively) reveal major changes upon ligand binding. Using the partial assignment of the methyl TROSY spectrum in the bound form, spectral effects during the GCDCA titration were classified qualitatively as follows and mapped on the corresponding crystal structure model (**c**, pdb 6E47, Nelson et al.): signals associated with slow exchange and consequently large chemical shift differences Δ*v* are indicated in red (as illustrated by the signal of bound Ile337, **d**), signals with intermediate, small and no shift differences are indicated in orange, yellow and grey, respectively (e.g. Val271, **e** and Val521, **f**).

### A model for MNV P-domain dimerization and GCDCA binding

Addition of GCDCA does not affect H^N^ cross peaks that are attributed to the monomeric P-domain. Thus, it is likely that GCDCA does not bind to P-domain monomers (cf. Fig. S8b, where the dimer peak “D” shifts whereas the corresponding monomer peak “M” just disappears). Accordingly, increasing GCDCA concentrations lead to depletion of monomers as the overall equilibrium is shifted towards the dimeric, bound state. This is supported by the disappearance of all monomer signals at equimolar concentrations of GCDCA, accounting for a majority of the observed complex spectral changes. Thus, for the following we assume that P-domain dimer formation must precede GCDCA binding. In the simplest binding model this leads to three coupled equilibria (Fig. 5) with two different dissociation constants. For the first equilibrium, the monomer-dimer exchange, we have derived a dissociation constant *K_D1_* from independent experiments as described above. From ab initio analysis of exchanging monomer and corresponding dimer cross peaks in concentration dependent methyl TROSY spectra we obtained a value of *K_D1_* of 13 μM (Fig. S5). This value is lower than the ones obtained from other experimental techniques and serves as a lower limit. Size exclusion chromatography provides an upper limit of 40 μM for *K_D1_* (Fig. 2c).

**Fig. 5.:**
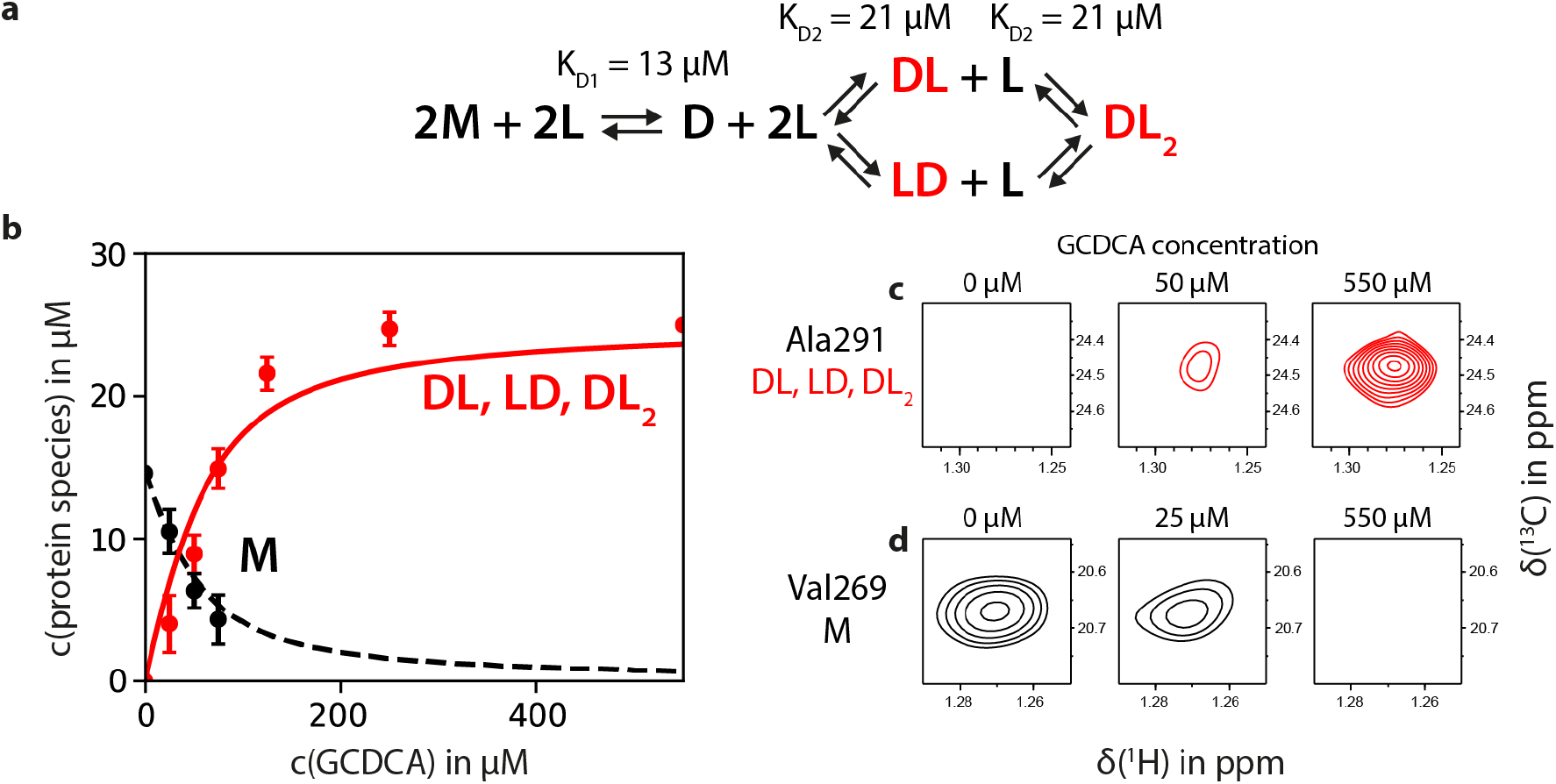
GCDCA titration data match a binding model of two consecutive, independent binding events to the dimeric protein only. **(a)** Equilibria linking monomers (M), dimers (D), GCDCA (L), and dimers with GCDCA bound (DL and DL_2_). **(b)** Intensity data of the methyl-TROSY-based GCDCA titration (cf. Fig. 3d) were fitted to a binding model in which the monomeric protein (M) is binding-incompetent and is depleted at higher ligand concentrations by removal of unbound dimer (D) through subsequent binding reactions. The dimerization dissociation constant *K_D1_* was determined independently by line shape analysis of methyl TROSY data (cf. Fig. S4). Concentrations of the bound species DL and DL_2_ were derived from signal intensities of slow exchange signals close to the binding pocket (**c**, indicated in red in Fig. 3c). Signals were averaged (n=11) and normalized to a protein concentration of 25 μM at saturation. Similarly, monomer signals (n=16) were normalized to a protein concentration in the absence of ligand as determined by *K_D1_* (**d**). Bound dimer data were fitted to the numerical solution to the system of differential equations describing this system as explained in detail in Fig. S10 to yield the dissociation constant of the ligand binding events, *K_D2_*. The theoretical curve (using the indicated *K_D_* values) is shown as a solid red line, monomer data was not used for fitting but is indicated for comparison in black. Error bars denote the standard deviation of the respective signal class before averaging.

Assuming that the two symmetrical GCDCA binding pockets of P-dimers are equal and independent, the second and the third equilibrium are characterized by a single microscopic dissociation constant *K_D2_*. Theoretical calculation of concentrations of monomers ([M]) and dimers ([DL], [LD], [DL2], cf. Fig. 5) using fixed values of *K_D2_* is possible by numerically solving eq. S7 for the concentration of monomers [M]. To illustrate the expected concentration curves during a titration of MNV P-domain with GCDCA concentrations of monomer and dimer species as a function of GCDCA concentrations were simulated and are shown in Fig. S10. We then monitored monomer and dimer cross peak intensities in methyl TROSY spectra through a titration of MNV P-domain with GCDCA. For determination of *K_D2_* we chose only well separated signals (e.g., red cross peak in Fig. 4d). We determined average cross peak intensities of dimer and monomer signals, resulting in corresponding experimental binding isotherms (Fig. 5b). Respective theoretical binding isotherms were obtained by systematically varying *K_D2_* and numerically solving eq. S7 for each value of *K_D2_*. Comparison of theoretical and experimental binding isotherms leads to a residual for each *K_D2_* value. The minimum of the residuals as a function of *K_D2_* represents the experimental *K_D2_* value (Fig. S11). The dissociation constant *K_D1_* was fixed during these calculations, either using the lowest or the highest independently determined values of 13 μM and 40 μM, respectively. This procedure furnishes a narrow window of values for the dissociation constant *K_D2_* around 20 μM.

### Dimer dissociation rate constants of murine and human norovirus P-domains differ by about four orders of magnitude

In order to put our results in a broader perspective we also studied the exchange between monomers and dimers for HuNoV P-dimers. We chose GII.4 Saga P-dimers that have been studied by NMR in our laboratory before (30, 38, 39). Recent native MS studies (40) have indicated the presence of monomeric species for GII.4 P-domains, posing the question of stability of P-dimers. In general, NMR allows examination of the thermodynamics and kinetics of such monomer-dimer equilibria at near-physiological conditions. In contrast to the native MS study our previous NMR experiments did not directly indicate the presence of traceable amounts of monomers, reflecting very stable P-dimers in solution. It follows that monomer-dimer exchange must be rather slow. In order to obtain the dissociation rate constant for GII.4 Saga P-dimers we employed analytical ion exchange chromatography (IEX), following a protocol established before to study spontaneous deamidation of a specific asparagine residue, Asn373, located in the histo blood group antigen (HGBA) binding pocket (30). Briefly, deamidation exclusively furnishes an iso-aspartate residue (iD) in position 373 and essentially switches off HBGA binding. We took advantage of the charge differences between the different dimeric species, termed “N/N” (both positions 373 are Asn), “iD/iD” (both positions 373 are iso-Asp), and “iD/N” (mixed) and employed analytical IEX to estimate the half-life for the deamidation process yielding a value of 1.6 days at 310 K (30). To study the dimerization equilibrium, we followed a similar strategy, employing GII.4 Saga P-domain point mutants. The point mutant N373Q does not undergo spontaneous deamidation and carries no negative charge in position 373. Purified fully deamidated iD/iD P-dimers also do not experience spontaneous deamidation and carry two extra negative charges compared to the N373Q mutant. Mixing N373Q P-dimers and iD/iD P-dimers leads to exchange of monomeric units over time yielding three types of P-dimers, homodimers iD/iD and Q/Q as well as mixed P-dimers iD/Q. The charge differences between iD/iD, Q/Q and iD/Q allow separation using analytical IEX (Fig. 6a) as described before (30). Starting with a 1:1 mixture of pure iD/iD and Q/Q P-dimers and subjecting aliquots of the mixture to quantitative IEX analysis at fixed time intervals over a period of 38 days yields the concentrations of homodimers and mixed dimers as a function of time (Fig. 6b). Assuming constant monomer concentrations throughout the experiment exchange of monomeric P-domains only depends on the rate constant *k_off_* for the dissociation of P-dimers (41). Fitting the concentration of mixed dimers to an exponential function yields a dissociation rate constant *k_off_* of 1.5 × 10^−6^ s^−1^, four orders of magnitude smaller than the corresponding value for MNV P-dimers. These results are supported by native mass spectrometry experiments performed on GII.4 Saga N/N P-dimers. The mass spectra almost exclusively show signals from P-dimers (Fig. 6c).

**Fig. 6:**
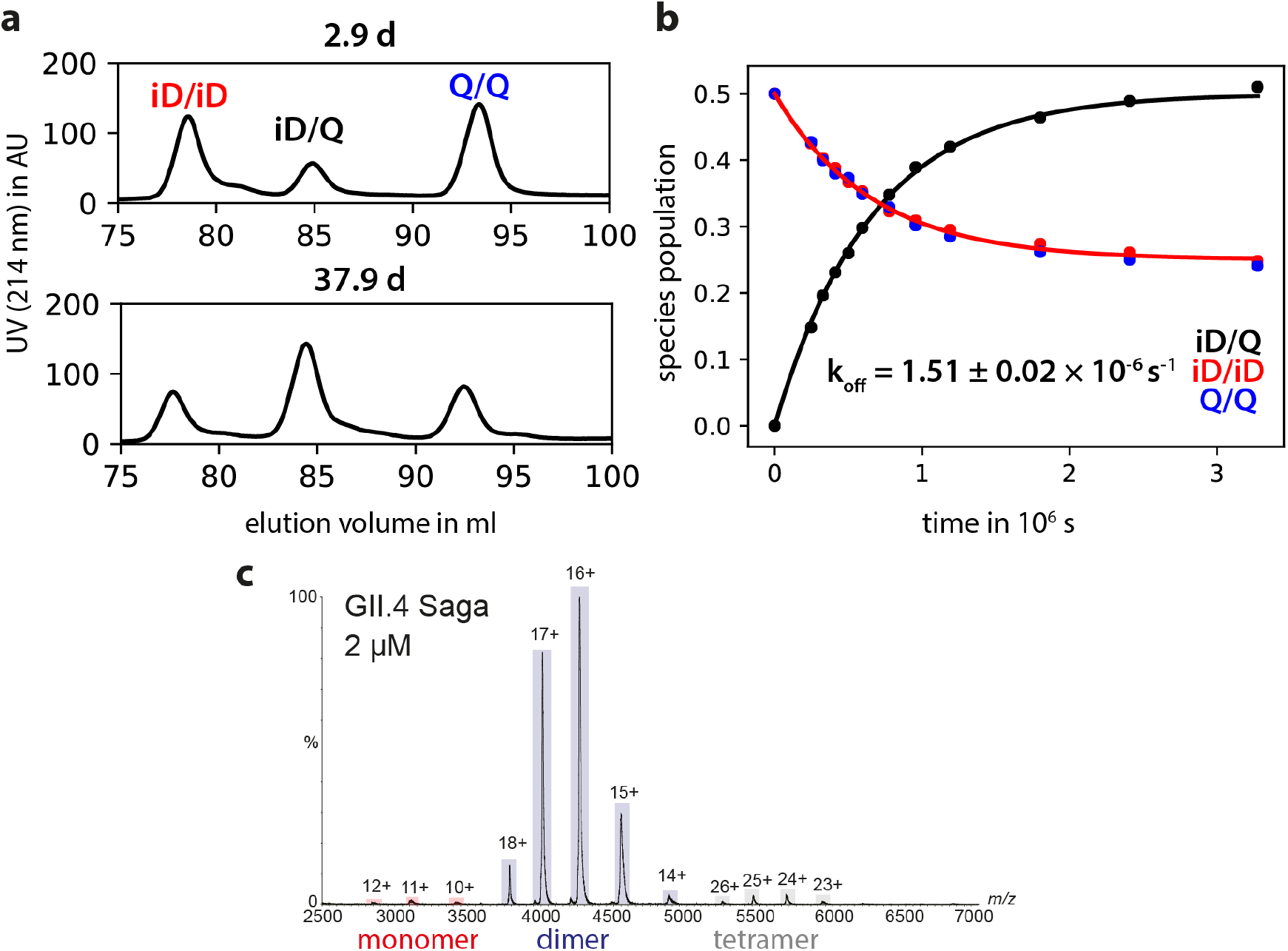
Dimer exchange kinetics of human NoV GII.4 Saga. Protruding domain samples which were deamidated at position 373 into a negatively charged iso-aspartate (iD) residue and point mutants with a neutral Gln at this position were mixed 1:1 and incubated at 298 K. The fraction of mixed 373 iD/Q dimers was quantified by analytical ion exchange chromatography at selected time intervals as explained in detail on the preceding page (**a**). Homo- and mixed dimer fractions reached the expected 1:2:1 ratio after approx. 40 d (**b**). Curve fitting yields a *k_off_* of 1.5 × 10^−6^ s^−1^. Native MS spectra of human GII.4 Saga N373 P-domain (N/N P-dimers) confirms mostly dimeric stoichiometry even at low protein concentration (**c**).

### Chemical shift perturbation NMR experiments reveal allosteric conformational changes that are sensed by a neutralizing antibody

Further analysis of CSPs of resonance signals in methyl TROSY spectra of MNV CW1 P-domains upon GCDCA binding (Fig. 4) relies on dissecting CSPs due to direct interactions of the ligand with amino acid side chain methyl groups located in the binding pocket from long-range effects observed for methyl groups located at positions distant to the binding site. The long-range effects reflect allosteric interactions upon GCDCA binding. It turns out that small long-range CSPs (Δ*v*_Eucl_ = 6 – 16 Hz) are observed almost throughout the P-domain reflecting allosteric interactions triggered by GCDCA binding (Fig. 4). In particular, we observe long-range CSPs of the order of 10 Hz (Δ*v*_Eucl_) for the methyl groups of the alanine residues 380, 381 and 382 located within the E’F’ loop (Fig. 7a-c). For Val378 flanking the E’F’ loop a significantly larger long-range effect is observed. As Val378 is one of the signals showing a large CSP (Δ*v*_Eucl_ ≫ 16 Hz) upon addition of GCDCA, and as we have no knowledge about the position of this signal in the apo form we can only state that the corresponding CSP must be larger than ca. 30 Hz (Δ*v*_Eucl_). It is important here to consider not only CSPs given as Euclidian distances between peak positions but to also analyze chemical shift changes in the ^1^H and ^13^C dimension individually. Changes of ^13^C chemical shifts of Ala methyl groups directly report on the backbone conformation (42, 43) and Ile, Leu, Val and Met methyl ^13^C chemical shifts carry information on the side chain rotamer populations (44–48). Separate ^1^H and ^13^C chemical shift perturbations are summarized in Table S2. It is interesting to note that for the Ala residues in the E’F’ loop it is almost exclusively the ^13^C CSPs that contribute to the overall effect. As the E’F’ loop is the region where neutralizing antibodies bind to the MNV P-domain we subjected MNV-1 particles to binding assays with a neutralizing antibody in the presence and absence of GCDCA or TCA. Here, we observed that the neutralizing MAbs 2D3 and 4F9 neutralize MNV-1 infection only in the absence of GCDCA, while TCA at indicated concentrations had no effect on immune escape, confirming our initial hypothesis (Fig. 7d).

**Fig. 7:**
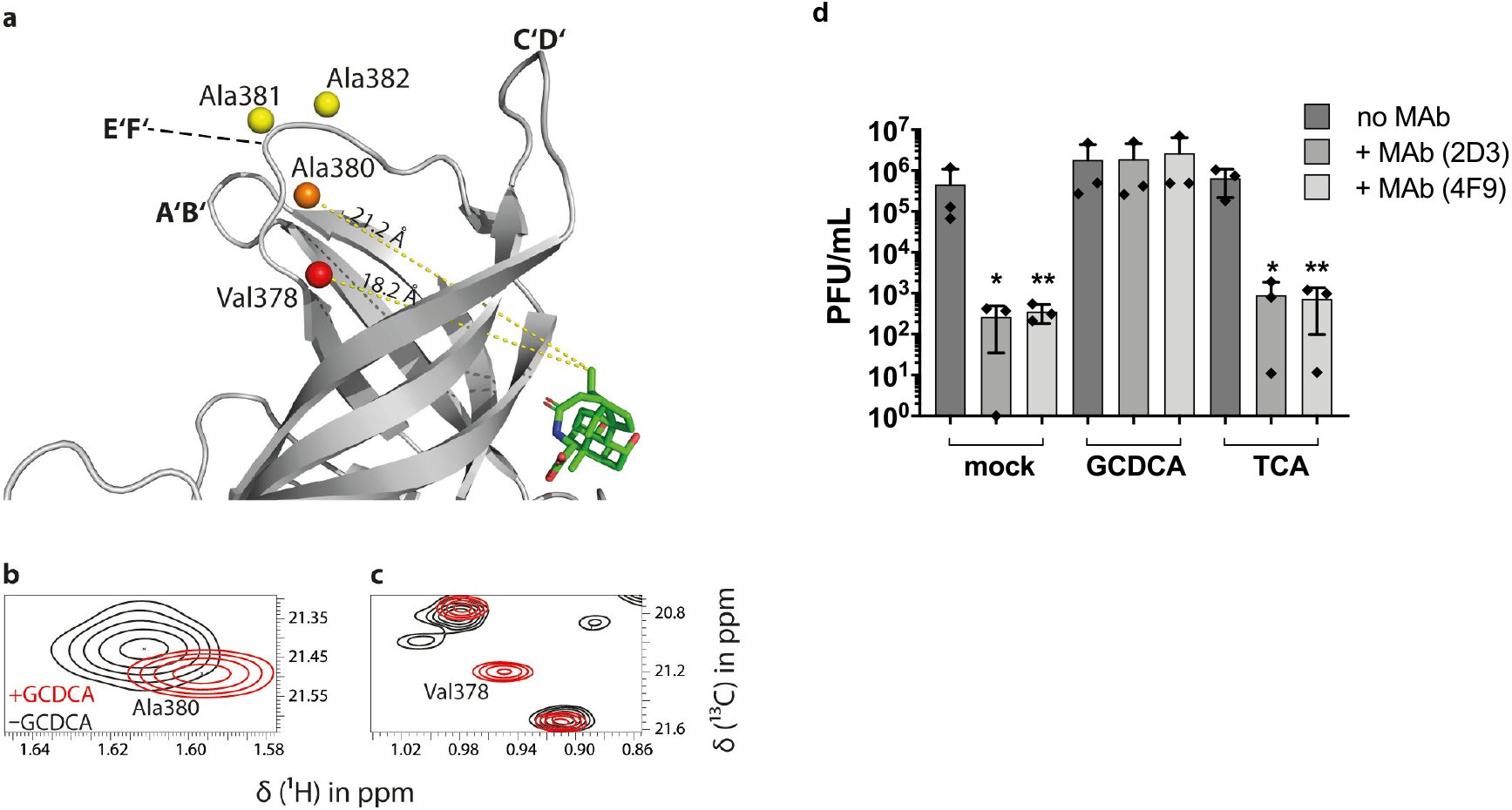
GCDCA induces long-range CSPs in the murine NoV P-domain E’F’ loop and causes escape from MAb recognition. **(a)** For the methyl group resonances of Ala380, Ala381, and Ala382 located in the E’F’ loop, and Val378 flanking this loop, long-range CSPs are observed in methyl TROSY spectra of MNV.CW1 P-domain (50 μM) in the presence of GCDCA. CSPs of methyl groups within the E’F’ loop (aa 378-388) plotted onto a crystal structure model (pdb 6E47, color code as in Fig. 4). These methyl groups are about 20 Å apart from the GCDCA binding pocket, reflecting an allosteric effect of GCDCA binding on the E’F’ loop. **(b, c)** The chemical shift perturbation with increasing amounts of GCDCA is exemplarily shown for Ala380 and Val378. At the highest GCDCA concentration of 550 μM, CSPs of 12 Hz, 6 Hz, and 9 Hz were measured for Ala380, Ala381, and Ala382, respectively. The Val378 cross peak **(c)** belongs to the class of “red signals” (Δ*v* ≫ 16 Hz, cf. Fig. 4d) and, therefore, the respective CSP must be significantly larger than ca. 30 Hz. The precise value cannot be reported since the apo-form peak has not been assigned yet. **(d)** MNV-1.CW3 immune escape from MAb 2D3 and 4F9 is inhibited by the bile acid GCDCA. MNV-1.CW3 was incubated with 1000 ng MAb (2D3 or 4F9) or without MAb (no MAb) in the absence of bile acid (mock), or in the presence of 500 μM GCDCA, or TCA. Plaque assays were performed in duplicate for at least three independent assays and plaque forming units (PFU) per ml were determined. Bars represent the mean (n=3) ± standard deviation (SD). Statistical analysis was performed using the unpaired *t*-test. The * indicates significance (*p*<0.05) compared to the respective treatment without MAb.

## Discussion

In the complete viral capsid, P-domains are connected with the underlying shell via an extended hinge comprising several amino acids. The possibility of loose instead of tight interactions between P-domains arranged as dimers in the viral capsid has not been considered before. Crystal structures of the P-domains feature dimers, insinuating that these dimers are “stable”. On the other hand, native MS data on HuNoV P-domains suggest that monomer-dimer equilibria play a role (40), and for MNV P-domains it has been noted that P-dimers may dissociate into monomers in solution (10). These observations prompted for a more detailed analysis of norovirus P-dimer dissociation in terms of thermodynamic stabilities and dissociation rates. We have used protein NMR spectroscopy supplemented by complementary biophysical techniques such as native MS, analytical IEX, and SEC to arrive at a comprehensive picture of P-dimer dissociation, revealing notable differences between human and murine noroviruses. Importantly, MNV P-domain dimer formation, in contrast to HuNoV, is strongly promoted by bile acid binding, at the same time triggering loop reorientations and subsequent immune escape. In the following we discuss our findings in the light of established knowledge.

A key finding of this study is that MNV P-domains co-exist as a mixture of dimers and monomers in aqueous solution under acidic conditions, which corresponds to the physiological environment in the gastrointestinal tract of the host (49). We have employed a simple binding model (Fig. 5a) to specify on- and off-rate constants (*k_on_* and *k_off_*) for P-dimer formation and dissociation, respectively, and a corresponding dissociation constant (*K_D1_*). The model also allows specification of a dissociation constant (*K_D2_*) for binding of GCDCA to P-dimers. The analysis of P-dimer dissociation is based on *ab initio* simulations (50) of concentration dependent methyl TROSY spectra of MNV.CW1 P-domains, and is substantiated by independent data from native MS and analytical size exclusion chromatography yielding values in a similar range (cf. Table 1 for a compilation of rate constants and dissociation constants). Notably, a dissociation rate constant *k_off_* of 0.04 s^−1^ translates into a dimer half-life *t*_1/2_ of ca. 17 s.

**Table 1.**
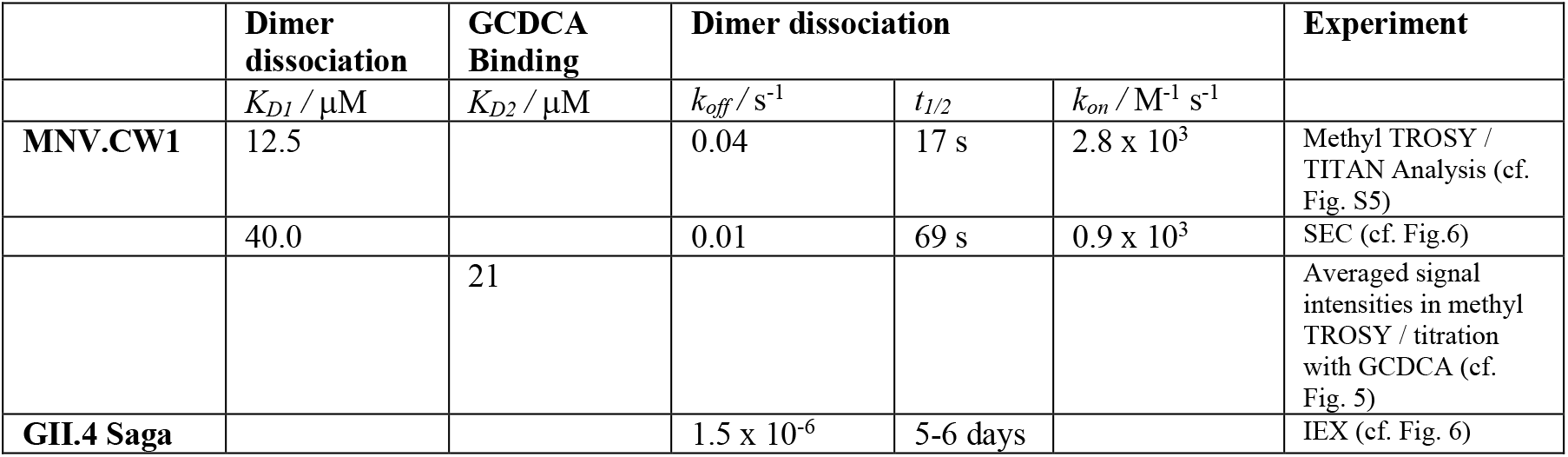
Dissociation constants and rate constants for MNV.CW1 and GII.4 Saga P-dimers.

Upon addition of a specific bile acid, GCDCA, the monomeric population disappears, reflecting substantial stabilization of P-domain dimers. As it is impossible to experimentally dissect the contributions of dimer stabilization and binding our model treats this certainly more complex process as one single step, yielding an apparent dissociation constant *K_D2_*. The model further assumes that monomeric P-domains are binding-incompetent, which is supported by CSP NMR experiments (cf. Figs. 4 and S9). Binding isotherms obtained from a methyl TROSY based CSP titration experiment with GCDCA (Fig. 5b) give no indication of cooperative effects between binding to the two symmetrical (10) GCDCA binding pockets. Therefore, the two sites are considered equal and independent. Based on these assumptions, the apparent dissociation constant *K_D2_* has been determined as described above with *K_D1_* kept fixed at either the smallest or the largest value determined independently (cf. Fig. S11). It turns out that *K_D2_* is not very sensitive to a variation of *K_D1_*, yielding a value for *K_D2_* of ca. 20 μM. This matches quite well with published data from an isothermal microcalorimetry (ITC) titration (10), where a value of 6 μM was reported for the dissociation constant. The difference may well be explained by the fact that in the ITC study a coupled equilibrium for P-dimer formation had not been accounted for.

In addition to promoting dimer formation GCDCA triggers remote (allosteric) conformational changes in the P-domain. In general, long-range CSPs are an indication of allosteric effects caused by ligand binding (37, 51). Here, GCDCA binding leads to a large number of long-range CSPs of methyl resonances in different regions of the P-domain, indicating respective conformational changes. It has been reported that the ^13^C chemical shifts of Ala methyl groups are correlated with the backbone dihedral angles ϕ and φ and, therefore, are sensitive reporters of conformational changes of the protein backbone upon, e.g., ligand binding (43). A systematic analysis of ^13^C CSPs of Ala methyl groups is currently limited since the fraction of assignments of corresponding chemical shifts in the apo form is still small. However, for the Ala residues located in the E’F’ loop, i.e., Ala residues 380, 381 and 382 an assignment was possible, and the resulting CSP values (Fig. 7 and Table S2) demonstrate that this loop undergoes an allosteric conformational change upon GCDCA binding. This is further supported by the observation that the chemical shift of the *proS* ^13^C methyl group of Val 378, which is flanking the E’F’ loop is strongly affected upon addition of GCDCA. All of these positions are about 20 Å away from the binding site (Fig. 7a). It has been described before that the transition between the open and the closed conformation is associated with reorientation of the A’B’, E’F’ and the C’D’ surface loops (18) in a concerted fashion. When bile salts bind to the P-dimer, the C’D’ loop moves up, pushing the E’F’ and the A’B’ loop into a conformation that is presumably not recognized by neutralizing antibodies (13, 19). The CSPs observed in the E’F’ loop represent direct experimental evidence for this hypothesis that so far has been missing. The A’B’ loop should also undergo conformational reorganization upon GCDCA binding but is lacking ^13^C-methyl reporter groups. Future work, e.g., based on measurements of residual ^13^C-^1^H dipolar couplings (52, 53) may complete the overall picture. The plasticity and conformational changes in the surface loops triggered by bile acid represent a novel allosteric immune escape pathway for murine noroviruses. It will now be interesting to test if respective escape mutations lead to similar conformational rearrangements.

As we have performed our studies with isolated P-domains the question arises whether the observed monomer-dimer equilibrium plays a role for native virus particles where the P-domains are linked to the S-domains. It is unlikely that MNV P-domain dimers completely dissociate when attached to the shell domain. However, it is entirely possible, that the relative movement of monomeric units associated with dimerization or dissociation plays a role for capsid plasticity. GCDCA binding causes the viral particle to change from an extended conformation to a contracted conformation. One can envision the P-domain having more degrees of freedom in the extended particle, where the P-domain is raised 10 – 15 Å above the shell. In particular, the two monomeric units may have enough translational degrees of freedom to partially dissociate generating a more “loose” overall conformational state. This would provide an explanation of the blurriness observed in cryo EM pictures (13) in the extended conformation due to the increased mobility at the lateral interphase. In the presence of GCDCA, when the P-domains are compacted onto the shell there is less motional freedom, which is further restricted by the fact that GCDCA stabilizes the dimeric form. Based on our data and on results published before by others (12, 13, 18, 19) we have devised a model reflecting the effects of GCDCA binding on MNV capsids and linking it to escape from antibody recognition (Fig. 8).

**Fig. 8.**
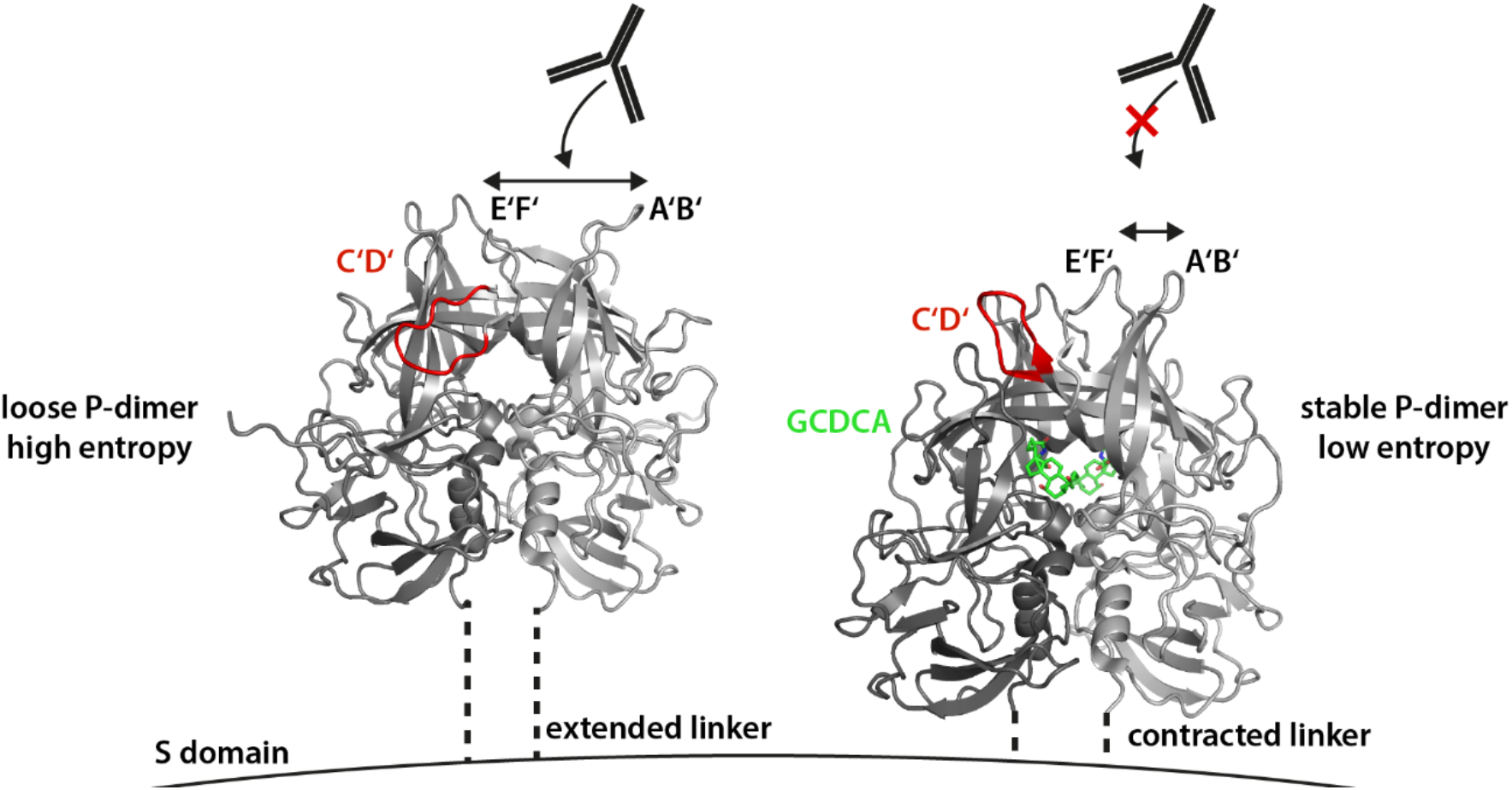
Model of immune escape and capsid contraction of MNV-1 P-dimers through binding of GCDCA. In the apo state (left), P-domains are elevated from the S domains and possess considerable mobility through partial dissociation of the P-dimer. Only this conformational state of the virion can be neutralized by the MAb 2D3. Tightening of P-domain interactions by GCDCA binding at the P-domain interface decreases conformational freedom and might entropically assist capsid contraction. The allosterically triggered rearrangement of the A’B’ and E’F’ loops prevents MAb binding to virions in the presence of bile acids.

The observed stabilization of P-dimers upon GCDCA binding also offers a new perspective for better understanding the contraction of the capsid. In the absence of GCDCA, P-domains protrude from the virus capsids, allowing several degrees of motional freedom. In particular, partial dissociation of P-dimers may add to conformational flexibility. Transition into a contracted, presumably more ordered conformation would be associated with a loss of conformational degrees of freedom and thus with a decrease in entropy. Therefore, in the absence of GCDCA contraction would be entropically unfavorable. As demonstrated in this study binding of GCDCA stabilizes P-dimers, eliminating the possibility of partial dissociation of P-dimers and, therefore, reducing the entropic penalty for capsid contraction.

The near-atomic cryo-EM structure of the full-length P-domain (residues 225-541) complexed with Fab A6.2 has recently been determined (T.J.S. et al., submitted for publication) and is consistent with these observations. The Fab is observed to bind to the P-domain in the “open” conformation and, from the structure of the complex, the Fab would not be able to bind to the bile-induced “closed” conformation. This was confirmed by binding studies showing that GCDCA decreases the affinity of the Fab binding to the P-domain. Further, comparisons between this P-domain dimer and the previous MNV capsid/GCDCA complex (54), shows that bile causes a rotation of the two subunits of the P-domain dimers that allows the P-domain to bind to the shell surface. Therefore, as shown here, the interactions between the two subunits in the P-domain dimer are remarkably plastic and play a pivotal role in the bile-induced transformation from a highly flexible expanded capsid to a more stable contracted state.

Interestingly, human norovirus P-domains are very different from MNV P-dimers with respect to the dimer-monomer equilibrium. The dissociation rate constant of GII.4 Saga P-dimers is about four orders of magnitude smaller than for MNV CW1 P-dimers, indicating a dissociation constant in the nano- to picomolar range. Moreover, HuNoV P-dimers provide no high-affinity binding pocket for GCDCA. Rather, GCDCA binding takes place with dissociation constants in the millimolar range (38). In the case of a GII.4 strain it could be shown that the respective binding pocket is located at the bottom of the P-dimers (38). For a rare genotype, GII.1, yet another bile acid binding site has been observed underneath the HBGA binding site (55).

Overall, dimer-monomer exchange has been identified as an additional cause for the plasticity of MNV capsids, offering a thermodynamically based explanation for capsid contraction upon GCDCA binding. Concomitant escape from recognition by a neutralizing antibody in the presence of GCDCA reflects allosterically induced conformational changes observed in the E’F’ loop. A picture emerges in which a small mediator molecule, a bile acid, triggers complex transformations in MNV but not in HuNoV capsids. Our study paves the way for biophysical and biological follow-up experiments to understand why MNV and HuNoV capsids behave so differently.

## Materials and Methods

### Protein biosynthesis

P-domain proteins were synthesized according to a previously published protocol (30). Briefly, *E. coli* BL21 DE3 cells were transformed with a plasmid containing genes for ampicillin resistance and a fusion protein of maltose-binding-protein (MBP) and the P-domain, separated by two His_8_-tags and a HRV3C cleavage site. Unlabeled protein was expressed in terrific broth medium, whereas [*U*-^2^H,^15^N] labeled P-domain was expressed in D_2_O-based minimal medium with deuterated glucose and ^15^NH_4_Cl (Deutero) as sole carbon and nitrogen sources, respectively. The isotopically labeled precursors for [*U*-^2^H], ɛ-[^13^C,^1^H_3_]-Met, δ1-[^13^C,^1^H_3_]-Ile, γ2-[^13^C,^1^H_3_]-Val, δ2-[^13^C,^1^H_3_]-Leu, β-[^13^C,^1^H_3_]-Ala-labeling of methyl groups (MIL^proS^V^proS^A -labeling) are given in Tab. S4. The precursors were dissolved in minimal medium (one fifth of the final culture volume) and added to the culture when an OD_600_ of 0.8 was reached. Cells were grown at 37 °C until an OD_600_ of 0.6-0.8 was reached. Then, expression was induced by addition of 1 mM IPTG for [*U*-^2^H,^15^N] labeled proteins and 0.1 mM IPTG for MIL^proS^V^proS^A-labeled proteins and growth was continued at 16 °C until the stationary phase was reached. Cells were lysed using a high-pressure homogenizer (Thermo). The lysate was clarified by ultra-centrifugation and subjected to Ni-NTA affinity chromatography to yield the pure fusion protein. The fusion protein was cleaved by addition of HRV3C protease and simultaneous dialysis against 20 mM sodium acetate buffer, 100 mM NaCl (pH 5.3). Ni-NTA affinity chromatography was repeated to separate the cleavage products. The P-domain protein was concentrated and applied to a preparative Superdex 16/600 200 pg size exclusion column (GE) using the aforementioned buffer as running buffer. Amino acid sequences of the cleaved P-domains are given in Tab. S1 of the supplement. Protein concentrations were quantified by UV spectroscopy using a molar extinction coefficient of 46870 M^−1^cm^−1^.

### Analytical size exclusion chromatography

Analytical size exclusion chromatography has been performed with a Superdex 75 Increase 3.2/300 column (GE Life Sciences) and 20 mM sodium acetate, 100 mM NaCl (pH 5.3) as running buffer with a flowrate of 0.075 ml min^−1^ at 278 K. To study the effect of ligand binding, 100 μM glycochenodeoxycholic acid (GCDCA, Sigma Aldrich) were added to the running buffer. Protein samples were applied with a 10 μl sample loop using a complete loop filling technique. UV absorption was monitored at 280 nm and 214 nm simultaneously. Apparent molecular weights as a function of feed protein concentration were fitted to the equation below (20) to yield an apparent dissociation constant *K_D_* of the protein-protein interaction as well as apparent molecular weights of the pure monomer and dimer.

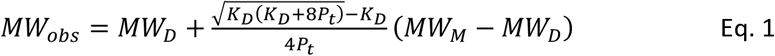

*MW_obs_* is the observed molecular weight according to the calibration of the column using standard proteins, *MW_D_* and *MW_M_* are the apparent molecular weights of the monomers and dimers, respectively, and *P_t_* is the total protein concentration applied onto the column.

A 1 ml HiTrap Butyl-HP column (GE) was used for hydrophobic interaction chromatography. Protein samples (240 and 270 μg ml^−1^ for pH and GCDCA data set, respectively) were prepared in different buffers and then subjected to isothermal denaturation at 45 °C. The effect of GCDCA binding was studied using 20-1000 μM GCDCA in 20 mM sodium acetate buffer, 100 mM NaCl (pH 5.3) with 30 min incubation time, while experiments on pH-dependence were performed in 75 mM sodium phosphate buffer, 100 mM NaCl with pH values ranging from 5.7 to 8 with 10 min incubation time. Prior to application onto the column 750 mM ammonium sulfate were added from a highly concentrated stock. The protein bound to the column in 20 mM sodium acetate, 750 mM ammonium sulfate (pH 5.3) and was eluted using a linear gradient over 5 column volumes up to 100 % of 20 mM sodium acetate (pH 5.3). A flow rate of 3 ml min^−1^ was used. The UV integral at 214 nm of a non-heat-treated control sample was normalized to 1. Experiments were performed as duplicates.

### Exchange kinetics of murine and human protruding domain dimers

P-domain monomers (“*M*”) and P-domain dimers (“*D*”) are assumed to exist in an exchange equilibrium (Eqs. 2–5) with *K_D_* being the dissociation constant (corresponds to *K_D1_* in Fig. 2), *P_t_* the total protein concentration, *k_on_* the association rate constant, *k_off_* the dissociation rate constant, and *k_ex_* the exchange rate constant:

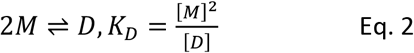

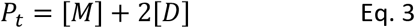

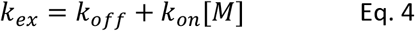

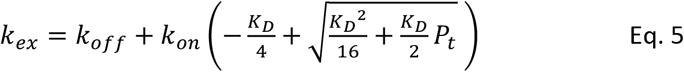

The dimer association rate *k_on_* can be approximated from the appearance of peak tailing at protein concentrations higher than 6 μM in the analytical size exclusion chromatography experiments (cf. Fig. 1c) using the relationship Eq. 6 derived in (56), which determines a minimum threshold value *N_k on_* for the observation of tailing:

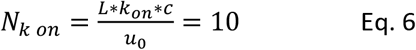

with *L* being the length of the column (30 cm), *c* being the lowest feed concentration at which tailing appears (6 μM), and *u_0_* being the linear flow rate (0.9325 cm min^−1^). As the determination of the lowest feed concentration at which tailing is observed is somewhat subjective we only give an approximate value for *k_on_*:

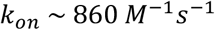

With *K_D_* = 12 μM and *P_t_* = 100 μM:

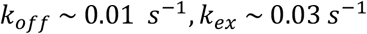

This is in good agreement with the upper limit for k_ex_ derived from the chemical shift difference of monomer and dimer signals at 100 μM sample concentration:

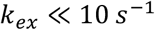

This is in stark contrast to the exchange kinetics of human norovirus GII.4 Saga protruding domain dimers that have exchange kinetics on the timescale of days, not seconds. GII.4 Saga P-domain proteins were synthesized as described elsewhere (30). We recently discovered an irreversible post-translational deamidation of Asn 373 into an iso-aspartate residue (iD). This reaction introduces a new negative charge into each monomer. The point mutant Asn373Gln does not undergo deamidation and carries no negative charge at this position. This charge difference of stable 373iD/iD and 373Q/Q homodimers can be exploited to analyze the kinetics of monomer exchange using analytical cation exchange chromatography (IEX, Fig. S3). Both types of dimers were mixed 1:1 with a total protein concentration of 1.24 mg ml^−1^ in 75 mM sodium phosphate buffer, 100 M NaCl, 0.02 % NaN_3_ (pH 7.3) and incubated at 298 K. At selected time points spanning 38 days aliquots were subjected to IEX experiments using the experimental conditions described in (30). With prolonged incubation times a new protein species can be observed corresponding to mixed dimers with an intermediate net charge. Exchange observed in such a mixing experiment only depends on *k_off_* as monomer concentrations are almost constant throughout the experiment (57).

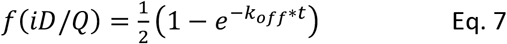

The fraction of mixed dimers observed at different time points has been fitted to the above equation to yield:

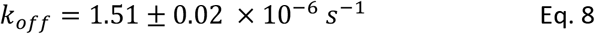

Extrapolation of these results to native GII.4 P-domains relies on the assumption that the type of the amino acid at position 373 does not change the dimerization equilibrium. At this point we would like to note that native MS experiments reproducibly reflect the presence of monomeric species for GII.4 Saga iD/iD P-dimers but not for the native, non-deamidated N/N P-dimers (40). Obviously, this is at odds with the NMR data giving no indication of different dimer stabilities of the two forms of the dimers. At present we have no conclusive explanation for these diverging observations, but we believe the very different experimental setup used in MS and NMR is responsible, suggesting that MS and NMR shed light on different, maybe related phenomena.

### NMR spectroscopy

NMR spectra were acquired at 298 K on either a 500 MHz Bruker Avance III or a 600 MHz Bruker Avance III HD NMR spectrometer with TCI cryogenic probes. Samples of [*U*-^2^H,^15^N]-labeled proteins were prepared in 20 mM sodium acetate buffer, 100 mM NaCl, 500 μM DSS-d_6_, and 10% D_2_O (pH* 5.3) unless stated otherwise. Methyl group labeled proteins were prepared in D_2_O containing 20 mM sodium acetate-d_3_, 100 mM NaCl, and 100 μM DSS-d_6_, (pH* 5.26).

NMR spectra were processed with TopSpin v3.6 (Bruker), signal intensities were obtained with CcpNmr Analysis v2.4.2 (58). Acquisition parameters are given in Tab. S3 or in the respective figure legends. Protein concentrations in NMR experiments concerning P-domain dimerization were 12.5-226 μM in ^1^H,^13^C HMQC-based experiments with methyl group labeled samples (600 MHz) and 24-200 μM in ^1^H,^15^N TROSY HSQC-based experiments (500 MHz). For GCDCA titration experiments, 100 μM [*U*-^2^H,^15^N]-labeled P-domain (500 MHz) and 50 μM MIL^proS^V^proS^A -labeled protein (600 MHz) were titrated separately with GCDCA from highly concentrated ligand stock solutions prepared in the respective sample buffers up to final concentrations of 300 μM and 550 μM, respectively. Signals in HMQC-based titration experiments were categorized according to the behavior of the slow exchange signals of unbound and bound protein species. Intensities of assigned, fully resolved signals of the bound species were used to obtain the GCDCA dissociation constant K_D2_. Intensities of these signals were normalized to the concentration of dimers at ligand saturation (25 μM) and then averaged. The system of equations Eq. S2-S5 describing a coupled equilibrium of dimerization and ligand binding of the dimer (cf. Fig. S10) was solved numerically with in-house Python (v2.7) scripts using SciPy’s (v1.2.1) *fsolve* function. *K_D2_* was derived from the averaged intensity data with a least-squares approach (cf. Fig. S11).

^15^N backbone relaxation data were obtained at 600 MHz using TROSY-based pulse schemes for measurement of *T_1_* and *T_1ρ_* relaxation times (59). Spectra were acquired with 128 increments in the indirect dimension and a relaxation delay of 3 s. Delays in the pulse sequence were 0 s, 0.36 s, 0.6 s, 1 s, and 2.48 s for determination of *T_1_* and 1 ms, 15 ms, 30 ms, and 50 ms for determination of *T_1ρ_*. Both experiments contained a spin-lock temperature compensation element of up to 50 ms. The spin-lock field strength *ω* in the *T_1ρ_* experiment was 1.4kHz, the carrier frequency in the ^15^N dimension was 117.5 ppm.

Rotational correlation times *τ_c_* of [*U*-^2^H,^15^N]-labeled murine NoV P-domains have been estimated using TRACT experiments at 500 MHz (36). The relaxation delay was set to 2 s. Experiments were measured for 80 min with 25 increasing delays in the pulse sequence of up to 0.2 s. Data were integrated from 6-9.5 ppm for murine NoV P-domains, normalized, and fitted to an exponential decay model for determination of average ^15^N *R_α_* and *R_β_* relaxation rates.

Assignment of methyl group signals in ^1^H,^13^C HMQC spectra was based on a 4D HMQC-NOESY-HMQC spectrum (60). The spectrum was acquired with a sample containing 500 μM protein and 700 μM GCDCA at 600 MHz. The protein was concentrated in Amicon centrifugal filters (10 KDa, Merck) in presence of 1:10 GCDCA. 30% non-uniform-sampling (NUS) were applied in the 4D spectrum with a Poisson gap schedule (61, 62) with 14087 NUS data points and a grid of 86 (^13^C) × 84 (^1^H) × 52 (^13^C) in the indirect dimensions. 512 data points were acquired in the direct dimension and the number of scans was set to 4 corresponding to a measurement time of approximately 7 d. The mixing time was 120 ms. The spectrum was reconstructed using multi-dimensional decomposition implemented in the TopSpin software (Bruker). Signals were first assigned to their corresponding amino acid type by synthesis of selectively MI-, IL^*proS*^V^*proS*^-L^*proS*^V^*proS*^-, V- and A-labeled samples. Expression media for selective V-labeling contained 80 mg/l leucine-d_10_ to suppress labeling of leucine methyl groups in presence of 2-^13^C-methyl-4-d_3_-acetolactate. NOEs between methyl group resonances of the MIL^proS^V^proS^A-labeled P-domain obtained from the 4D HMQC-NOESY-HMQC experiment gave information about short- and long-range spatial restraints of the corresponding methyl groups. This information was compared to an existing crystal structure model (pdb 6e47) for assignment of HMQC signals to their corresponding methyl groups (32, 39).

Line shape analysis of ^1^H,^13^C HMQC spectra with TITAN (35) was performed with MIL^proS^V^proS^A - labeled samples at different protein concentration as stated above using TITAN’s built-in dimerization model. Spectra were processed in NMRPipe (63) prior to analysis. First, positions and linewidths of seven isolated putative monomer peaks were fitted using spectra of 12.5 μM and 25 μM P-domain. Then, positions and linewidths of the seven corresponding dimer peaks in spectra with 226 μM and 100 μM P-domain were fitted accordingly. Finally, peak positions of monomer and dimer peaks were held constant and all linewidths, the dissociation rate k_off_ and K_D1_ were fitted using the dimerization binding model at all six P-domain concentrations with the linewidths for monomer and dimer signals derived above as starting values. Parameter uncertainties were obtained by bootstrap error analysis.

### Mass spectrometry

Proteins were subjected to buffer exchange to 150 mM ammonium acetate, pH 5.3 (murine P-domains) or pH 7.3 (GII.4 Saga P-domain) at 4 °C via Micro Bio-Spin 6 columns (Bio-Rad) according to the manufacturer’s protocol. Native MS measurements were performed using 2 to 92 μM purified P domains. Mass spectra were acquired at room temperature (25 °C) in positive ion mode on an LCT mass spectrometer modified for high mass (Waters, UK and MS Vision, the Netherlands) with a nano-electrospray ionization source. Gold-coated electrospray capillaries were produced in house for direct sample infusion. Capillary and sample cone voltages were 1.30 kV and 200 V, respectively. The pusher was set to 100-150 μs. Pressures were 7 mbar in the source region and 6.2 × 10^−2^ to 6.5 × 10^−2^ mbar argon in the hexapole region. A spectrum of a 25 mg/mL cesium iodide solution from the same day was applied for calibration of raw data using the MassLynx software (Waters, UK).

### Determination of dissociation constants from MS

Monomer and dimer peak areas were summed over all corresponding charge states. The relative dimer peak area was calculated and plotted against the total protein concentration (*P_t_*). The *K_D_* was then determined by global non-linear least squares fitting of Eq. 9 (64) to the dataset using OriginPro 2016 software:

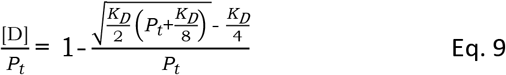

### Cell culture

Murine microglia cells (BV-2) were maintained in Dulbecco’s Modified Eagle Medium (Gibco) supplemented with 5% FCS (C-C-Pro), 2 mM L-glutamine (Biozym), 0.1 mM non-essential amino acids (Biozym), and 100 units/ml penicillin and streptomycin (Biozym) (DMEM-5) as described (Hwang 2014). Cells were incubated at 37°C with 5% CO_2_ and 95% humidity.

Hybridoma suspension cell lines producing 2D3 and 4F9 antibodies were a kind gift from Christiane Wobus (University of Michigan, USA). Briefly, cells were maintained in Iscove’s Modified Dulbecco’s Medium (IMDM, Life Technologies) supplemented with 10% FCS (C-C-Pro), 0.5 mM L-glutamine (Biozym), and 100 units/ml penicillin and streptomycin (Biozym) (IMDM-10). Cells were cultivated in spinner flasks at 40 rpm at 37°C with 5% CO_2_ and 95% humidity with loosened lids for gas-exchange. Passaging was performed replacing 50% of the ‘old’ conditioned medium with 50% fresh medium diluting the cells to a concentration of at least 1 × 10^5^ cells per ml.

### Purification of anti-MNV antibodies

MAb against MNV (2D3 and 4F9) were produced from hybridoma suspension cultures. As MAb 2D3 and 4F9 are IgA antibodies, they cannot be purified by protein A/G chromatography and were purified by size-exclusion chromatography. Briefly, cultures were grown to a density of at least 1 × 10°6 cells per ml and pelleted by centrifugation (10 min 1000 rpm). Cell culture medium replaced with serum free medium (Capricorn Scientific Hybridoma Plus) supplemented with 100 units/ml penicillin and streptomycin (Biozym) maintaining a cell density of 1 × 10^6^ cells per ml. Cultures were incubated in spinner flasks at 37°C with 5% CO_2_ for 3 days. Suspensions were clear-centrifuged (30 min 3000 rpm 4°C) and the cell pellet was discarded. The supernatant containing the antibodies was sterile-filtered (0.2 μM) and purified by size-exclusion chromatography equilibrated with PBS. Purity was assessed by SDS-PAGE and Coomassie staining and protein concentration was determined using the BCA protein assay kit (Pierce).

### Virus Stocks

MNV-1.CW3 (GV/MNV1/2002/USA) was cultivated in BV-2 cells and used at passage 6. Stocks were produced in BV-2 cells in cell culture medium in the absence of serum (DMEM-0) as described previously (Wegener 2018).

### Virus quantification by plaque assay

Briefly, BV-2 cells were seeded in a six well plate at a density of 5 × 10^5^ cells per well and incubated over night at 37°C with 5% CO_2_ and 95% humidity. Monolayers were inoculated (37°C, 5% CO_2_, 1h) with 500 μl virus stock serially diluted in DMEM10 and swirled ca. every 15 min to facilitate even virus adsorption. The inoculum was removed, and cells were covered with 2 ml overlay medium containing 0.6% Avicel RC-591 in DMEM supplemented with 10% (v/v) FCS (C-C-Pro), 20 mM HEPES (Biozym), 4 mM L-glutamine (Biozym), 0.2 mM non-essential amino acids (Biozym), and 200 units/ml penicillin and streptomycin (Biozym) and incubated (37°C, 5% CO_2_, 48h). After 48h, the overlay medium was gently aspirated, and cells were washed once with phosphate buffered saline pH 7.4 (PBS). Monolayers were then stained with staining solution (PBS with 0.1 mg/ml (w/v) erythrosine B). After 15 – 20 min incubation at room temperature, the staining solution was aspirated. Plaque assays were generally performed in duplicate and plaque-forming units (PFU) per ml were determined.

### Plaque neutralization assay

The plaque neutralization assay is based on the plaque assay described above. Briefly, 1 × 10^6^ PFU of MNV-1.CW3 were diluted in PBS in the presence or absence of 1 μg of IgA MAb MAb 2D3 or 4F9 and in the presence or absence of 500 μM bile acid (sodium glycochenodeoxycholate GCDCA, or sodium taurocholate hydrate TCA, Sigma Aldrich) followed by enumeration by plaque assay. Plaque neutralization assays were generally performed in duplicate and plaque-forming units (PFU) per ml were determined.

### Thermal stability assay

The thermal stability assay is based on the plaque assay described above. Briefly, 1 × 10^7^ PFU per ml of MNV-1.CW3 were incubated at temperatures ranging between 0 and 56°C for 2 or 6 h in the presence or absence of 500 μM bile acid (sodium glycochenodeoxycholate GCDCA, Sigma Aldrich) by enumeration by plaque assay.

### Statistics

Differences in plaque -forming units (PFU) are presented as means ± standard deviation (SD) of duplicate samples from at least three independent experiments. Statistical analysis was performed using a two-tailed *t* test with two degrees of freedom. All statistical analyses were carried out using the Prism software package (GraphPad Software, CA).

## Acknowledgments

This research was funded by the *Deutsche Forschungsgemeinschaft* (DFG) via grants Pe494/12-2 (T.P.) and TA1093-2 (S.T.) within the research unit FOR2327 (ViroCarb). T.P. thanks the *State of Schleswig-Holstein* for supplying the NMR infrastructure (European Funds for Regional Development, LPW-E/1.1.2/857). C.F. thanks the *Studienstiftung des deutschen Volkes* for a fellowship. J.D. and C.U. acknowledge funding from FOR2327 ViroCarb. C.U. acknowledges funding from the Leibniz Association through SAW-2014-HPI-4 grant. The Heinrich-Pette-Institute, Leibniz Institute for Experimental Virology is supported by the Free and Hanseatic City of Hamburg and the Federal Ministry of Health. T.J.S. acknowledges funding from the NIH, grant 1R01-AI141465.

## Notes

### Competing Interest Statement

The authors have declared no competing interest.

## References

1. J. van Beek et al., Molecular surveillance of norovirus, 2005–16: an epidemiological analysis of data collected from the NoroNet network. The Lancet Infectious Diseases 18, 545–553 (2018).

2. M. D. Kirk et al., World Health Organization Estimates of the Global and Regional Disease Burden of 22 Foodborne Bacterial, Protozoal, and Viral Diseases, 2010: A Data Synthesis. PLoS Med 12, e1001921 (2015).

3. M. de Graaf, J. van Beek, M. P. Koopmans, Human norovirus transmission and evolution in a changing world. Nat Rev Microbiol 14, 421–433 (2016).

4. M. K. Jones et al., Enteric bacteria promote human and mouse norovirus infection of B cells. Science 346, 755–759 (2014).

5. K. Ettayebi et al., Replication of human noroviruses in stem cell–derived human enteroids. Science 353, 1387–1393 (2016).

6. S. Taube et al., A mouse model for human norovirus. mBio 4, e00450–00413 (2013).

7. J. W. Perry, S. Taube, C. E. Wobus, Murine norovirus-1 entry into permissive macrophages and dendritic cells is pH-independent. Virus Res 143, 125–129 (2009).

8. R. O. Orchard et al., Discovery of a proteinaceous cellular receptor for a norovirus. Science 353, 933–936 (2016).

9. K. Murakami et al., Bile acids and ceramide overcome the entry restriction for GII.3 human norovirus replication in human intestinal enteroids. Proc Natl Acad Sci U S A 10.1073/pnas.1910138117 (2020).

10. C. A. Nelson et al., Structural basis for murine norovirus engagement of bile acids and the CD300lf receptor. Proc Natl Acad Sci U S A 10.1073/pnas.1805797115 (2018).

11. M. L. Mallory, L. C. Lindesmith, P. D. Brewer-Jensen, R. L. Graham, R. S. Baric, Bile Facilitates Human Norovirus Interactions with Diverse Histoblood Group Antigens, Compensating for Capsid Microvariation Observed in 2016-2017 GII.2 Strains. Viruses 12(2020).

12. H. Q. Smith, T. J. Smith, The Dynamic Capsid Structures of the Noroviruses. Viruses 11 (2019).

13. M. B. Sherman et al., Bile Salts Alter the Mouse Norovirus Capsid Conformation: Possible Implications for Cell Attachment and Immune Evasion. Journal of virology 93, e00970–00919 (2019).

14. J. Jung et al., High-resolution cryo-EM structures of outbreak strain human norovirus shells reveal size variations. Proc Natl Acad Sci U S A 10.1073/pnas.1903562116 (2019).

15. J. S. Snowden et al., Dynamics in the murine norovirus capsid revealed by high-resolution cryo-EM. PLoS Biol 18, e3000649 (2020).

16. U. Katpally, C. E. Wobus, K. Dryden, H. W. t. Virgin, T. J. Smith, Structure of antibody-neutralized murine norovirus and unexpected differences from viruslike particles. J Virol 82, 2079–2088 (2008).

17. C. Song et al., Dynamic rotation of the protruding domain enhances the infectivity of norovirus. PLoS Pathog 16, e1008619 (2020).

18. S. Taube et al., High-resolution x-ray structure and functional analysis of the murine norovirus 1 capsid protein protruding domain. Journal of virology 84, 5695–5705 (2010).

19. A. O. Kolawole et al., Norovirus Escape from Broadly Neutralizing Antibodies Is Limited to Allostery-Like Mechanisms. mSphere 2(2017).

20. C. M. Yu, S. Mun, N. H. Wang, Theoretical analysis of the effects of reversible dimerization in size exclusion chromatography. J Chromatogr A 1132, 99–108 (2006).

21. A. Shortland et al., Pathology caused by persistent murine norovirus infection. J Gen Virol 95, 413–422 (2014).

22. T. Kilic, A. Koromyslova, V. Malak, G. S. Hansman, Atomic structure of the murine norovirus protruding domain and sCD300lf receptor complex. J Virol 10.1128/JVI.00413-18 (2018).

23. S. Schütz, R. Sprangers, Methyl TROSY spectroscopy: A versatile NMR approach to study challenging biological systems. Progress in nuclear magnetic resonance spectroscopy 10.1016/j.pnmrs.2019.09.004 (2019).

24. C. Gobl, T. Madl, B. Simon, M. Sattler, NMR approaches for structural analysis of multidomain proteins and complexes in solution. Prog Nucl Magn Reson Spectrosc 80, 26–63 (2014).

25. D. P. Frueh, Practical aspects of NMR signal assignment in larger and challenging proteins. Prog Nucl Magn Reson Spectrosc 78, 47–75 (2014).

26. Y. Xu, S. Matthews, TROSY NMR spectroscopy of large soluble proteins. Topics in current chemistry 335, 97–119 (2013).

27. V. Tugarinov, V. Kanelis, L. E. Kay, Isotope labeling strategies for the study of high-molecular-weight proteins by solution NMR spectroscopy. Nat Protoc 1, 749–754 (2006).

28. V. Tugarinov, L. E. Kay, An Isotope Labeling Strategy for Methyl TROSY Spectroscopy. Journal of Biomolecular NMR 28, 165–172 (2004).

29. U. Katpally et al., High-resolution cryo-electron microscopy structures of murine norovirus 1 and rabbit hemorrhagic disease virus reveal marked flexibility in the receptor binding domains. J Virol 84, 5836–5841 (2010).

30. A. Mallagaray et al., A post-translational modification of human Norovirus capsid protein attenuates glycan binding. Nat Commun 10, 1320 (2019).

31. A. Mallagaray, J. Lockhauserbäumer, G. S. Hansman, C. Uetrecht, T. Peters, Attachment of Norovirus to Histo Blood Group Antigens: A Cooperative Multistep Process. Angew Chem Int Ed 54, 12014–12019 (2015).

32. A. Proudfoot, A. O. Frank, F. Ruggiu, M. Mamo, A. Lingel, Facilitating unambiguous NMR assignments and enabling higher probe density through selective labeling of all methyl containing amino acids. J Biomol NMR 65, 15–27 (2016).

33. V. Tugarinov, L. E. Kay, I. Ibraghimov, V. Y. Orekhov, High-Resolution Four-Dimensional 1H-13C NOE Spectroscopy using Methyl-TROSY, Sparse Data Acquisition, and Multidimensional Decomposition. Journal of the American Chemical Society 127, 2767–2775 (2005).

34. A. D. Bain, Chemical exchange in NMR. Progress in nuclear magnetic resonance spectroscopy 43, 63–103 (2003).

35. C. A. Waudby, A. Ramos, L. D. Cabrita, J. Christodoulou, Two-Dimensional NMR Lineshape Analysis. Sci Rep 6, 24826 (2016).

36. D. Lee, C. Hilty, G. Wider, K. Wüthrich, Effective rotational correlation times of proteins from NMR relaxation interference. Journal of magnetic resonance 178, 72–76 (2006).

37. L. Shi, L. E. Kay, Tracing an allosteric pathway regulating the activity of the HslV protease. Proc Natl Acad Sci U S A 111, 2140–2145 (2014).

38. R. Creutznacher et al., Chemical-Shift Perturbations Reflect Bile Acid Binding to Norovirus Coat Protein: Recognition Comes in Different Flavors. Chembiochem 21, 1007–1021 (2020).

39. C. Muller-Hermes, R. Creutznacher, A. Mallagaray, Complete assignment of Ala, Ile, Leu(ProS), Met and Val(ProS) methyl groups of the protruding domain from human norovirus GII.4 Saga. Biomol NMR Assign 10.1007/s12104-020-09932-z (2020).

40. J. Dülfer et al., Glycan-induced protein dynamics in human norovirus P dimers depend on virus strain and deamidation status. bioRxiv 10.1101/2020.10.07.329623, 2020.2010.2007.329623 (2020).

41. B. R. Capraro et al., Kinetics of endophilin N-BAR domain dimerization and membrane interactions. J Biol Chem 288, 12533–12543 (2013).

42. K. Pederson et al., NMR characterization of HtpG, the E. coli Hsp90, using sparse labeling with (13)C-methyl alanine. J Biomol NMR 68, 225–236 (2017).

43. R. Godoy-Ruiz, C. Guo, V. Tugarinov, Alanine Methyl Groups as NMR Probes of Molecular Structure and Dynamics in High-Molecular-Weight Proteins. Journal of the American Chemical Society 132, 18340–18350 (2010).

44. L. Siemons et al., Determining isoleucine side-chain rotamer-sampling in proteins from (13)C chemical shift. Chem Commun (Camb) 55, 14107–14110 (2019).

45. D. F. Hansen, L. E. Kay, Determining valine side-chain rotamer conformations in proteins from methyl 13C chemical shifts: application to the 360 kDa half-proteasome. J Am Chem Soc 133, 8272–8281 (2011).

46. D. F. Hansen, P. Neudecker, L. E. Kay, Determination of Isoleucine Side-Chain Conformations in Ground and Excited States of Proteins from Chemical Shifts. Journal of the American Chemical Society 132, 7589–7591 (2010).

47. G. L. Butterfoss et al., Conformational dependence of 13C shielding and coupling constants for methionine methyl groups. Journal of Biomolecular NMR 48, 31–47 (2010).

48. F. A. Mulder, Leucine side-chain conformation and dynamics in proteins from 13C NMR chemical shifts. Chembiochem 10, 1477–1479 (2009).

49. E. L. McConnell, A. W. Basit, S. Murdan, Measurements of rat and mouse gastrointestinal pH, fluid and lymphoid tissue, and implications for in-vivo experiments. J Pharm Pharmacol 60, 63–70 (2008).

50. C. A. Waudby, T. Frenkiel, J. Christodoulou, Cross-Peaks in Simple Two-Dimensional NMR Experiments from Chemical Exchange of Transverse Magnetisation. Angewandte Chemie 131, 8876–8880 (2019).

51. G. P. Lisi, J. P. Loria, Solution NMR Spectroscopy for the Study of Enzyme Allostery. Chem Rev 10.1021/acs.chemrev.5b00541 (2016).

52. R. V. Williams, J. Y. Yang, K. W. Moremen, I. J. Amster, J. H. Prestegard, Measurement of residual dipolar couplings in methyl groups via carbon detection. J Biomol NMR 73, 191–198 (2019).

53. R. Sprangers, L. E. Kay, Probing Supramolecular Structure from Measurement of Methyl 1H-13C Residual Dipolar Couplings. J Am Chem Soc 129, 12668–12669 (2007).

54. M. B. Sherman et al., Bile salts alter the mouse norovirus capsid conformation; possible implications for cell attachment and immune evasion. Journal of Virology 12, e00970–00919 (2019).

55. T. Kilic, A. Koromyslova, G. S. Hansman, Structural Basis for Human Norovirus Capsid Binding to Bile Acids. Journal of virology 93, e01581–01518 (2019).

56. C.-M. Yu, S. Mun, N.-H. L. Wang, Theoretical analysis of the effects of reversible dimerization in size exclusion chromatography. Journal of Chromatography A 1132, 99–108 (2006).

57. B. R. Capraro et al., Kinetics of endophilin N-BAR domain dimerization and membrane interactions. The Journal of biological chemistry 288, 12533–12543 (2013).

58. W. F. Vranken et al., The CCPN data model for NMR spectroscopy: development of a software pipeline. Proteins 59, 687–696 (2005).

59. N. A. Lakomek, J. Ying, A. Bax, Measurement of (1)(5)N relaxation rates in perdeuterated proteins by TROSY-based methods. J Biomol NMR 53, 209–221 (2012).

60. J. Wen, P. Zhou, J. Wu, Efficient acquisition of high-resolution 4-D diagonal-suppressed methyl-methyl NOESY for large proteins. Journal of magnetic resonance 218, 128–132 (2012).

61. S. G. Hyberts, H. Arthanari, S. A. Robson, G. Wagner, Perspectives in magnetic resonance: NMR in the post-FFT era. J Magn Reson 241, 60–73 (2014).

62. S. G. Hyberts, S. A. Robson, G. Wagner, Exploring signal-to-noise ratio and sensitivity in non-uniformly sampled multi-dimensional NMR spectra. J Biomol NMR 55, 167–178 (2013).

63. F. Delaglio et al., NMRPipe: A multidimensional spectral processing system based on UNIX pipes. Journal of Biomolecular NMR 6, 277–293 (1995).

64. C. Thiele, W. B. Huttner, The disulfide-bonded loop of chromogranins, which is essential for sorting to secretory granules, mediates homodimerization. J. Biol. Chem. 273, 1223–1231 (1998).

