## Supplementary Figures and Tables for "Murine norovirus capsid plasticity – Glycochenodeoxycholic acid stabilizes P-domain dimers and triggers escape from antibody recognition"

### SI Appendix

##### This PDF file includes:

Figs. S1 to S11

Tables S1 to S4

References (1-3)

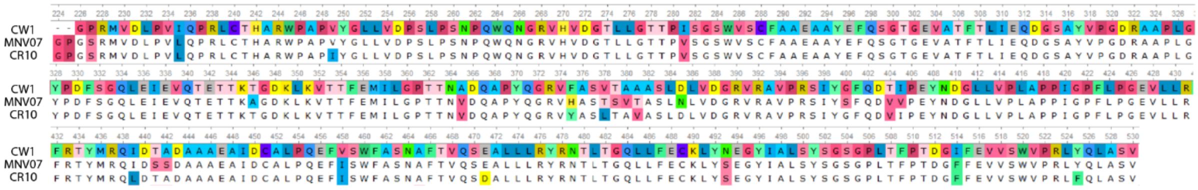

**Fig. S1**: Sequence alignment of recombinant CW1, MNV07 and CR10 P-domains from MNV used in this study. The sequences correspond to the GenBank entries given in Tab. S1.

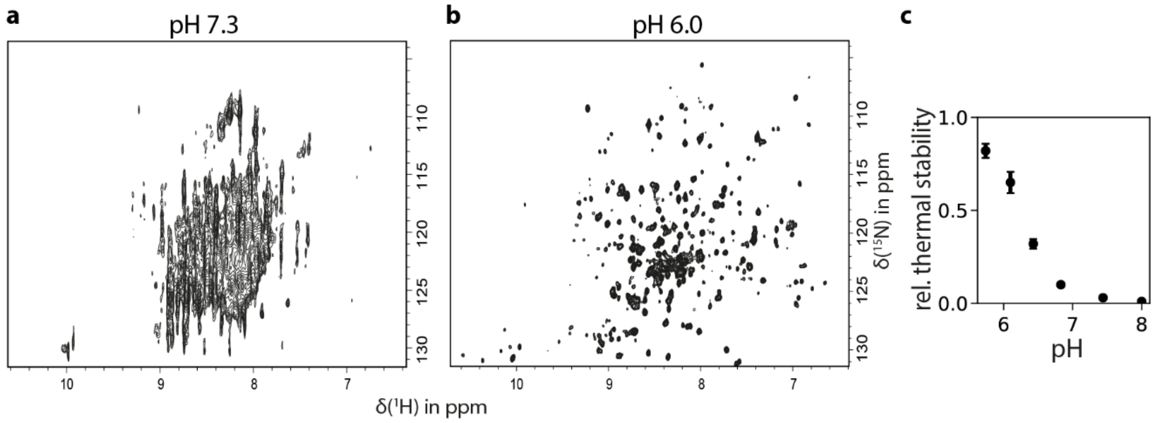


**Fig. S2**: **pH-dependent stability of murine NoV P-domains**. Purification of MNV P-domains in buffers at pH 7.3 leads to irreversible protein unfolding and aggregation (**a**). A TROSY HSQC spectrum of [$U$-$^{2}$H,$^{15}$N] labeled MNV07 P-domains in 20 mM sodium phosphate buffer (pH* 7.3) shows little chemical shift dispersion and strong line broadening. In contrast, samples in 20 mM sodium phosphate buffer, 100 mM NaCl (pH* 6) give spectra with dispersed, sharp NH signals indicating a well-ordered and stable protein (**b**). The thermal stability of MNV P-domains decreases with increasing pH (**c**). P-domains from the strain CR10 were subjected to isothermal denaturation in 75 mM sodium phosphate buffer, 100 mM NaCl at different pH values at 45 °C and subsequent hydrophobic interaction chromatography (HIC). The UV absorption in HIC experiments can be used to quantify the amount of non-denatured protein. UV integrals were normalized against a non-heat-treated control. HIC experiments were performed as duplicates, the respective percentage of deviation is given as error bars. The spectra in **a** and **b** were acquired with 15 µM protein concentration and 672 scans, and 30 µM and 136 scans, respectively. Additionally, spectrum **a** was acquired with 128 increments in the indirect dimension. Other acquisition parameters are given in Tab. S2.

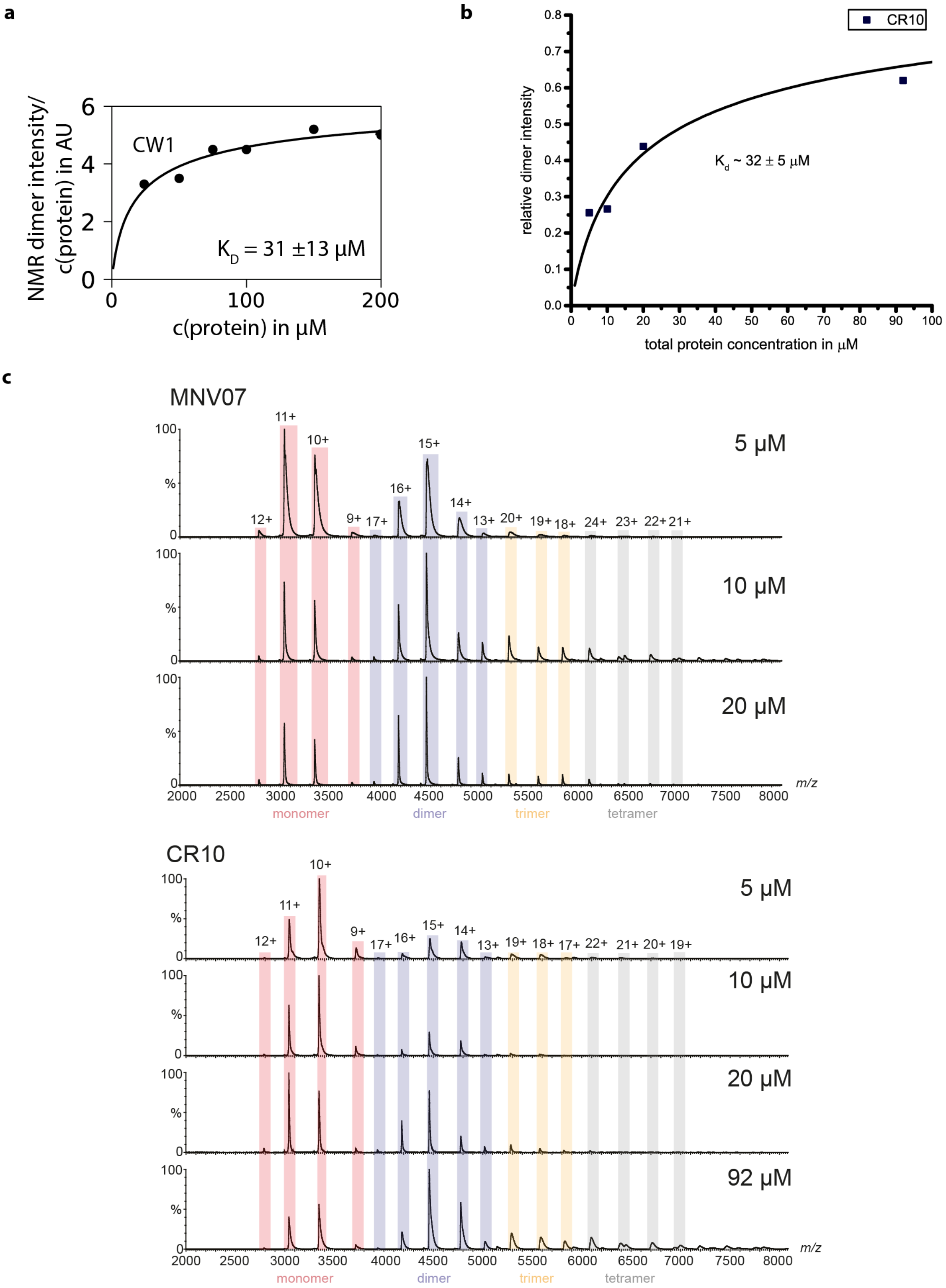


**Fig. S3**: **The MNV P-domain has a weak tendency to form dimers**. (a) The dissociation constant of the dimerization reaction was estimated based on the signal intensity of dimer signals (cf. Fig. 2b) in $^{1}$H,$^{15}$N TROSY HSQC spectra of [$U$-$^{2}$H,$^{15}$N] labeled MNV CW1 P-domains at different total protein concentrations. The signal intensities of 9 dimer signals (6.9|123.8 ppm, 8.5|130 ppm, 10.0|130.2 ppm, 8.8|125.6 ppm, 8.6|126 ppm, 7.5|131 ppm, 8.8|128.9 ppm, 9|112 ppm, 8.1|128.6 ppm) were averaged and fitted against the law of mass action (Eq. S1, below). Monomer-dimer ratios at different protein concentrations as determined by native MS (c) were fitted to the law of mass action to yield the dimerization dissociation constant for the MNV strain CR10 (b).

a

b

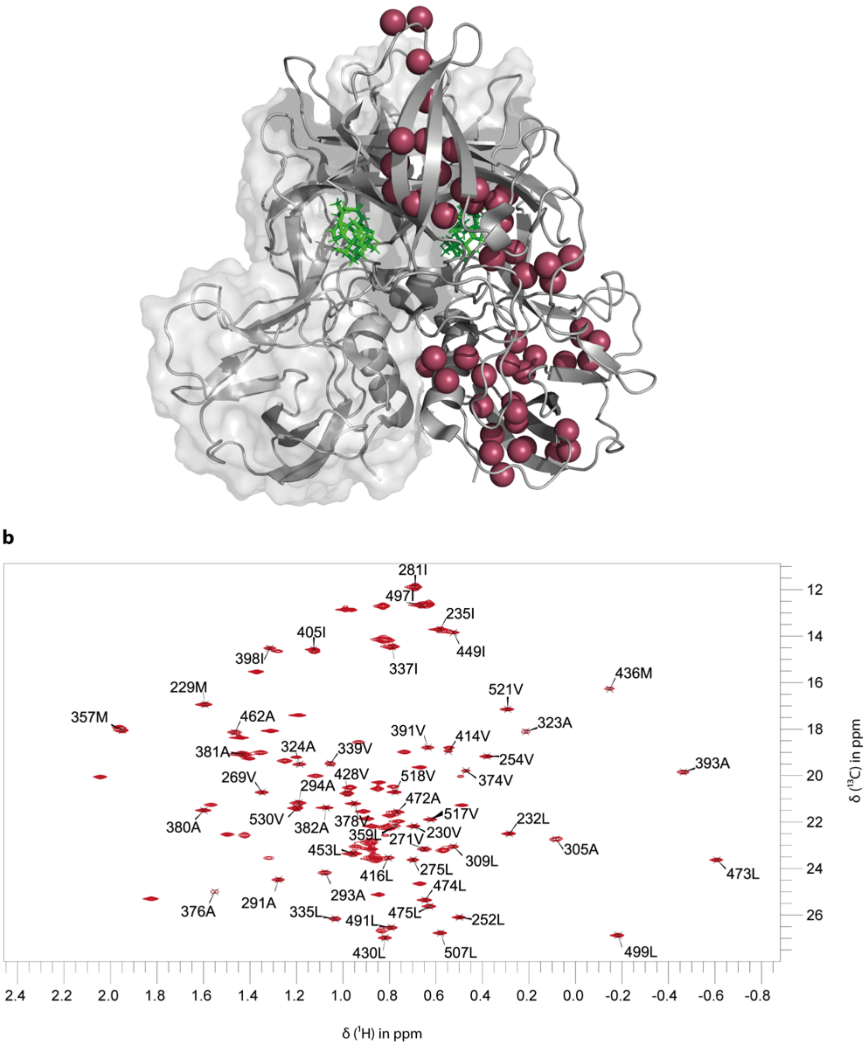


**Fig. S4**: Partial assignment of MNV CW1 MIL$^{ProS}$V$^{ProS}$A-labeled P-domain $^{1}$H,$^{13}$C HMQC resonances. (**a**) MNV P-Domain crystal structure (pdb 6e47). Methyl groups corresponding to assigned resonances are highlighted as colored spheres and are distributed over the entire protein. (**b**) $^{1}$H,$^{13}$C HMQC spectrum of 500 µM MIL$^{ProS}$V$^{ProS}$A-labelled MNV P-domain in presence of 700 µM GCDCA measured at 600 MHz. Assigned resonances are denoted with a cross and their corresponding amino acid type and number.

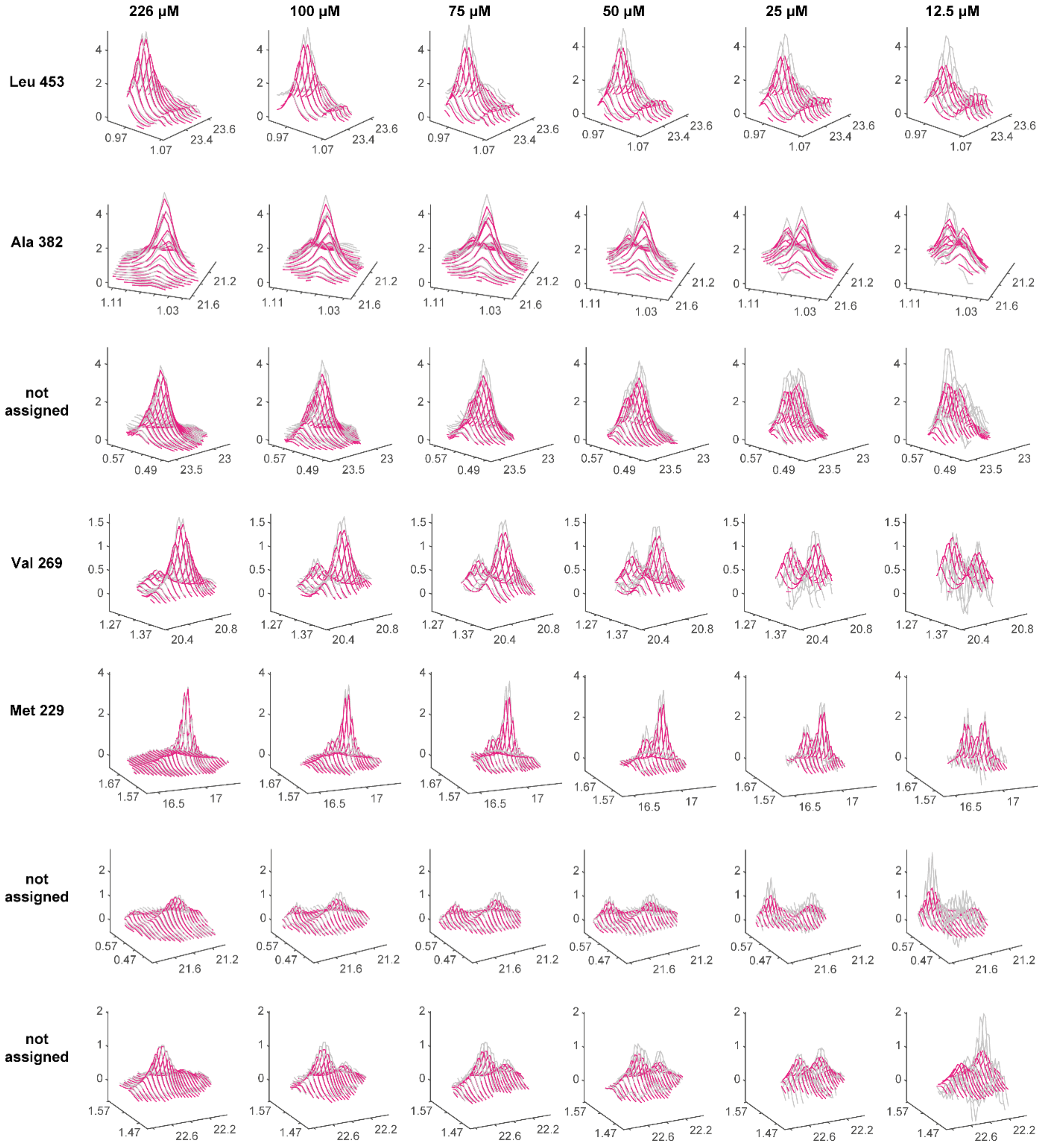


**Fig. S5**: 2D line shape analysis of $^{1}$H,$^{13}$C HMQC spectra of MIL$^{proS}$V$^{proS}$A-labeled samples at different protein concentrations using TITAN (1). Experimental and fitted spectra of several monomer-dimer signal pairs are shown in grey and magenta, respectively. This approach results in a dimerization $K_{D1}$ of 12 ± 1.5 µM and a dissociation rate $k_{off}$ of 0.035 ± 0.066 $s^{-1}$.

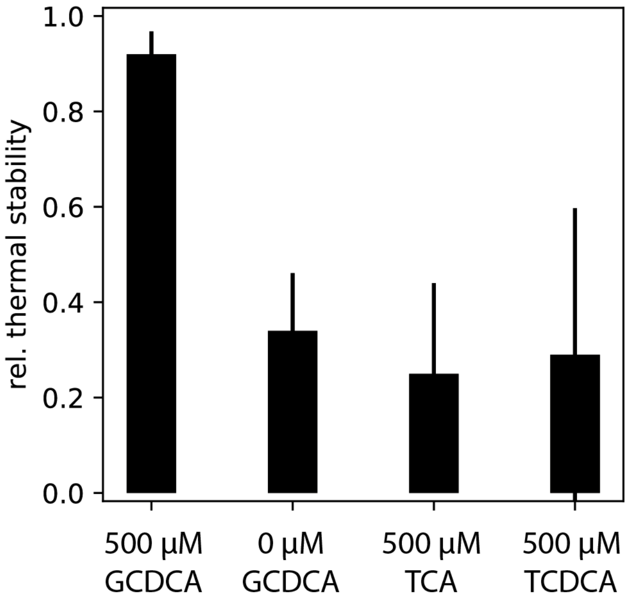


**Fig. S6**: **Increase of MNV P-domain thermal stability by ligand binding is specific for GCDCA**. Isothermal denaturation was performed for 30 min at 45 °C and analysis of non-denatured P-domain via HIC was performed by integration of UV absorption at 214 nm. GCDCA = glycochenodeoxycholic acid, TCA = taurocholic acid, TCDCA = taurochenodeoxycholic acid. HIC experiments were performed as duplicates, the respective percentage of deviation is given as error bars.

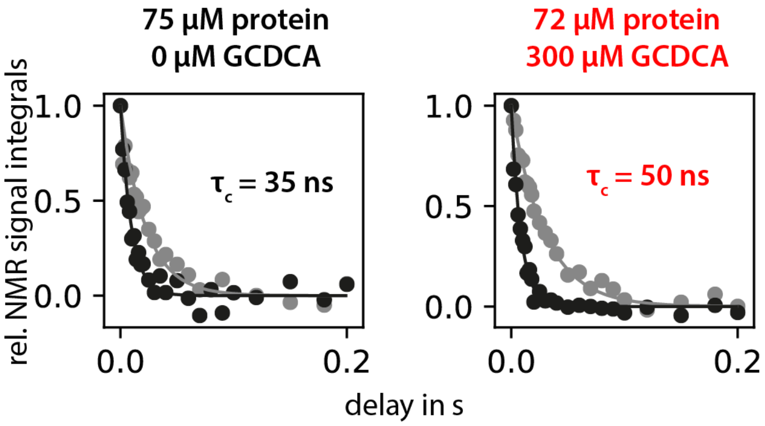


**Fig. S7: Rotational correlation time of MNV P-domains increases upon GCDCA addition**. TRACT NMR experiments can be used to determine rotational correlation times $\tau_c$ by comparison of the relaxation behavior of the different transitions (black and grey curves, respectively) associated with the protein's $^{15}N$ amide nuclei (2). The data shows an increase in correlation time upon GCDCA binding due to shift of equilibrium towards dimers from a sample containing a mixture of monomers and dimers in the absence of GCDCA (cf. Fig 2b). Data were acquired at 298 K on a 500 MHz spectrometer in 20 mM sodium acetate buffer, 100 mM NaCl (pH 5.3).

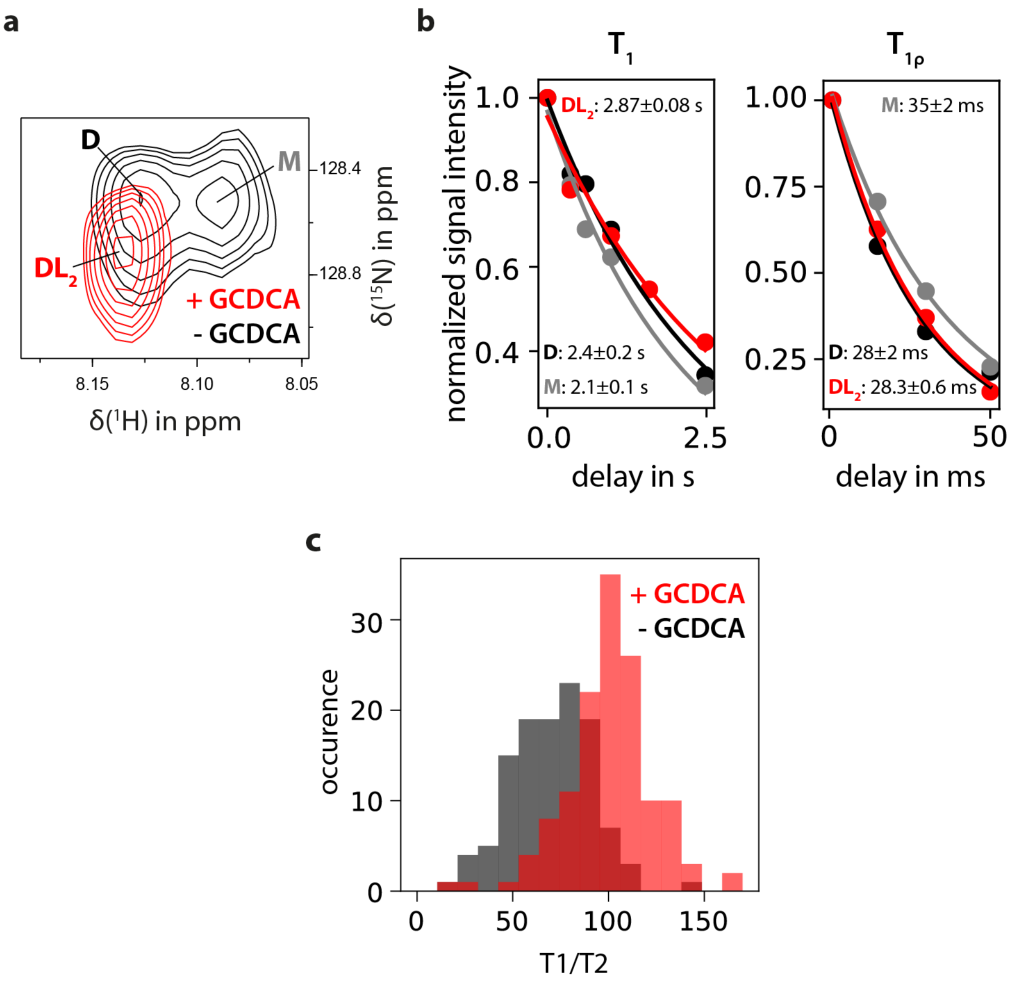


**Fig. S8: Changes in NMR relaxation times indicate altered protein dynamics upon dimerization and ligand binding**. $^{15}$N $T_1$ and $T_{1\rho}$ relaxation times (3) can be measured for individual amino acid signals in $^{1}$H,$^{15}$N TROSY HSQC spectra of MNV CW1 samples with and without GCDCA (**a**). For some spectral regions, signals can be assigned to monomeric (M), dimeric (D), or ligand-bound, dimeric ($DL_2$) protein species. Relaxation times can be obtained by curve fitting of decaying signal intensities with increasing delays during which relaxation can occur. Relaxation times were found to vary between the different protein species (**b**), e.g. between the signals corresponding to the different states of one amino acid shown in (**a**). The ratio $T_1/T_2$ is sensitive towards protein dynamics on the ps-ns time scale. Global analysis of $T_1/T_2$ for a high number of amino acids in the absence and in the presence of GCDCA (n=116 and 134, respectively) reveals pronounced differences between the unbound and bound protein states (**c**). $^{15}$N $T_1$ and $T_{1\rho}$ relaxation experiments were measured with a sample containing 230 µM [*U*-$^{2}$H,$^{15}$N]-labeled MNV1 P-domain in the absence of GCDCA and with a sample containing 150 µM protein and 500 µM GCDCA. Samples were prepared in 20 mM sodium acetate, 100 mM NaCl (pH 5.3) and contained 10 % $D_2O$.

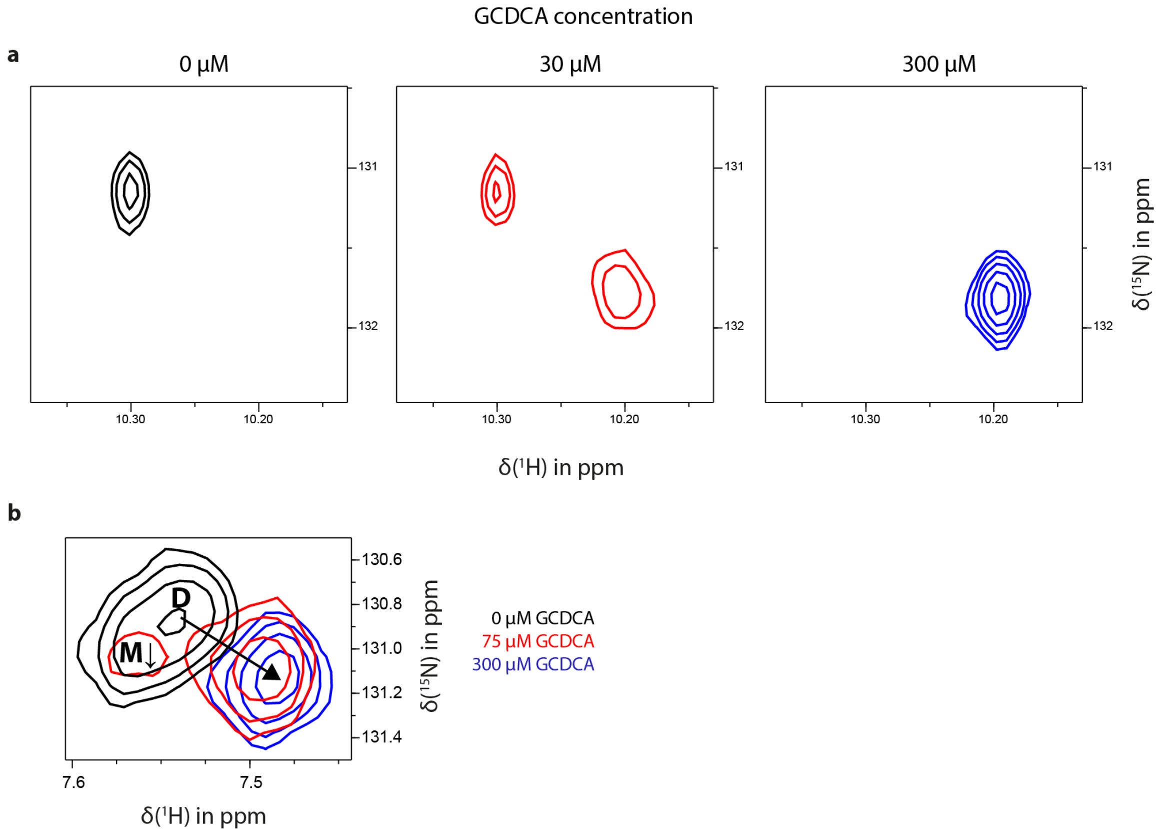


**Fig. S9**: Representative sections from $^{1}H,^{15}N$ TROSY HSQC spectra of 100 µM [$U$-$^{2}H,^{15}N$] labeled MNV CW1 P-domain with increasing GCDCA concentrations show both signals in slow exchange (**a**, Δν = 60 Hz) and fast exchange (**b**, Δν = 30 Hz). Signals attributable to the monomer (M) disappear with increasing GCDCA concentrations, whereas some dimer signals (D) display chemical shift perturbations (CSP). Samples were prepared in 20 mM sodium acetate buffer, 100 mM NaCl (pH 5.3).

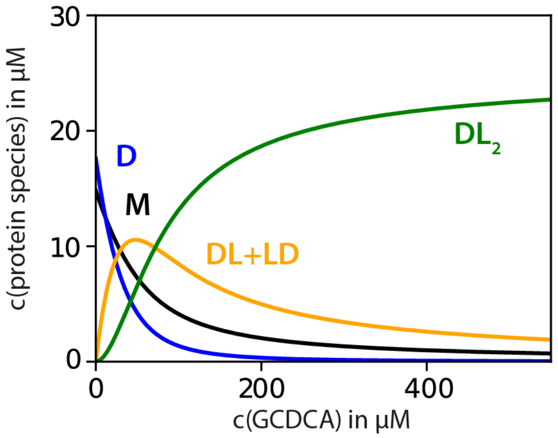


**Fig. S10: Predictions from a binding model of two consecutive, independent binding events to the dimeric protein only**. The evolution of the different protein species with increasing concentrations of the ligand GCDCA was simulated using the numerical solution of Eq. S7 below. $M$ = monomers, $D$ = dimers, $DL$ and $LD$ = dimer with one bound ligand, $DL_2$ = dimer with two bound ligand molecules. Model parameters were as follows: $K_{D1}$ = 12.5 µM, $K_{D2}$ = 21 µM, total protein concentration 50 µM.

**Equations describing coupled equilibria of dimerization and consecutive, microscopic ligand-binding reactions with independent, identical binding pockets with monomers incapable of binding**

$$2M + 2L \rightleftharpoons D + 2L \rightleftharpoons DL + L \rightleftharpoons DL_2$$

$$K_{D1} = \frac{[M]^2}{[D]}, K_{D2} = \frac{[D][L]}{[DL]} = \frac{[D][L]}{[LD]} = \frac{[LD][L]}{[DL_2]} = \frac{[DL][L]}{[DL_2]} \qquad (\text{Eq. S2} - \text{S3})$$

with $DL$ and $LD$ denoting the two dimer species with one bound ligand molecule that be distinguished microscopically.

$$[L]_t = [L] + [DL] + [LD] + 2[DL_2] \qquad (\text{Eq. S4})$$

$$[M]_t = [M] + 2[D] + 2[DL] + 2[LD] + 2[DL_2] \qquad (\text{Eq. S5})$$

With $[DL]$ and $[LD]$ being always equal, rearranging equations S2-S3 and substituting $[DL]$, $[DL_2]$ in ligand mass balance (Eq. S4) leads to:

$$0 = 2\frac{[D]}{{K_{D2}}^2}[L]^2 + \left(1 + \frac{2[D]}{K_{D2}}\right)[L] - [L]_t$$

Substituting $[D]$ with dimerization term:

$$[L] = -\left(\frac{K_{D1}{K_{D2}}^2}{4[M]^2} + \frac{K_{D2}}{2}\right) + \sqrt{\left(\frac{K_{D1}{K_{D2}}^2}{4[M]^2} + \frac{K_{D2}}{2}\right)^2 + \frac{[L]_t K_{D1}{K_{D2}}^2}{2[M]^2}} \qquad (\text{Eq. S6})$$

Substituting $[D]$, $[DL]$, $[LD]$, and $[DL_2]$ in protein mass balance (Eq. S5):

$$0 = [M] + \frac{2[M]^2}{K_{D1}} + \frac{4[M]^2}{K_{D1}K_{D2}}[L] + \frac{2[M]^2}{K_{D1}{K_{D2}}^2}[L]^2 - [M]_t \qquad (\text{Eq. S7})$$

This equation can be solved numerically for $[M]$ after inserting Eq. S6 as detailed in the materials and methods.

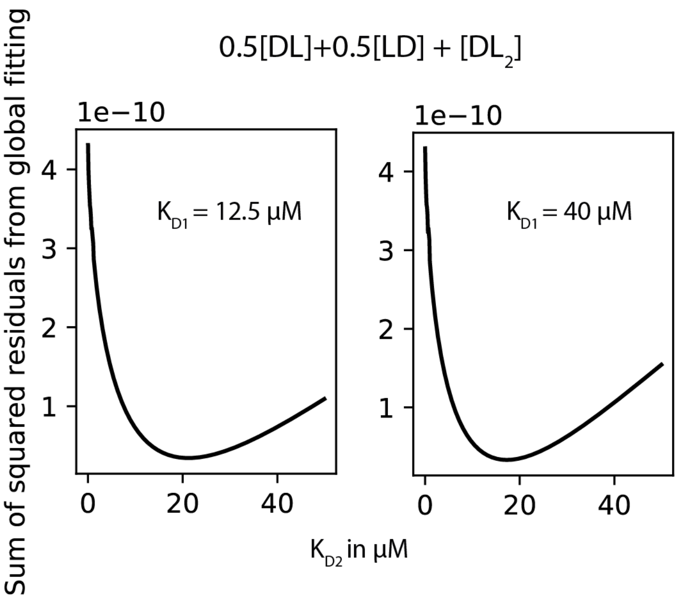


**Fig. S11**: Residuals from fitting of slow exchange signals in methyl TROSY spectra (cf. Fig. 3d and Fig. 4) during a GCDCA titration to the binding model given in Fig 5, given $K_{D1}$ = 12.5 µM (left, cf. Fig. S4) or $K_{D1}$ = 40 µM (cf. Fig. 1c). Data were fitted to the numerical solution of Eq. S7. The protein species *DL* and *LD* contribute only half to the total intensity of the slow exchange signal as only one monomeric subunit is bound. The data analysis process is detailed in the legend of Fig. 5.

**Tab. S1**: Amino acid sequences of used P-domain proteins. Additional N-terminal amino acids have been added to create the enzymatic cleavage site.

| Strain | GenBank ID | Additional N-terminal aa |
|---|---|---|
| CW1 | aa 228-530 of entry DQ285629 | GP |
| MNV07 | aa 228-530 of entry AET79296 | GPGS |
| CR10 | aa 228-530 of entry ABU55613 | GPGS |
| Saga2006 | aa 225-530 of entry AB447457 | GPGS |

**Tab. S2**: Alanine chemical shifts and chemical shift perturbations (CSPs) in the absence and in the presence of saturating amounts of GCDCA. Some signal positions in the apo state could not be determined due to slow exchange of the bound and unbound forms and CSPs that were too large to transfer the assignment from the bound state.

| **Ala** | **position apo $^{1}$H [ppm]** | **position apo $^{13}$C [ppm]** | **position GCDCA-saturated $^{1}$H [ppm]** | **position GCDCA-saturated $^{13}$C [ppm]** | **Euclidean CSP [Hz]** | **CSP in $^{13}$C [Hz]** |
|---|---|---|---|---|---|---|
| 291 | 1.214 | 24.589 | 1.276 | 24.474 | 39.6 | 14.3 |
| 293 | n.d. | n.d. | 1.077 | 24.176 | n.d. | n.d. |
| 294 | n.d. | n.d. | 1.189 | 21.187 | n.d. | n.d. |
| 305 | 0.083 | 22.379 | 0.081 | 22.714 | 41.8 | 41.8 |
| 323 | 0.219 | 18.041 | 0.213 | 18.061 | 4.7 | 2.5 |
| 324 | 1.159 | 19.466 | 1.184 | 19.508 | 15.9 | 5.2 |
| 380 | 1.610 | 21.419 | 1.597 | 21.492 | 12.3 | 9.1 |
| 381 | 1.428 | 19.140 | 1.428 | 19.093 | 5.9 | 5.9 |
| 382 | 1.071 | 21.316 | 1.071 | 21.385 | 8.6 | 8.6 |
| 393 | -0.333 | 20.267 | -0.466 | 19.836 | 96.7 | 53.9 |
| 462 | 1.457 | 18.211 | 1.464 | 18.125 | 11.7 | 10.8 |
| 472 | 0.759 | 21.600 | 0.767 | 21.563 | 6.6 | 4.6 |

**Tab. S3**: Acquisition parameters of 2D NMR-based pulse programs unless stated otherwise in the respective figure legends. TD denotes the number of increments in the respective dimensions, O1 is the center of the spectral window, SW is the sweep width, AQ is the acquisition time and NS is the number of scans.

| Experiment | Field strength | F2 (TD, O1, SW, AQ) | F1 (TD, O1, SW, AQ) | NS |
|---|---|---|---|---|
| $^{1}H,^{15}N$ TROSY HSQC | 500 MHz | 2048<br>4.7 ppm<br>16 ppm<br>128 ms | 256<br>117.5 ppm<br>35 ppm<br>72 ms | 32 |
| $^{1}H,^{13}C$ HMQC (meTROSY) | 600 MHz | 512<br>0.8 ppm<br>3.5 ppm<br>122 ms | 256<br>17 ppm<br>18 ppm<br>47 ms | 8 |

**Tab. S4**: Final concentrations of precursors for MILVA-labeling of MNV-P-domains.

| Amino acid | Precursor | Concentration |
|---|---|---|
| $L^{ProS}$ , $V^{ProS}$ | 2-$^{13}$C-methyl-4-d$_3$-acetolactate | 195 mg/L |
| I | 2-ketobutyricacid-4-$^{13}$C-3, 3-d$_2$ | 72 mg/L |
| A | L-alanine-$^{13}$C-d$_2$<br>succinate-d$_4$ | 0.6 g/L<br>3.75 g/L |
| M | L-methionine-(methyl-$^{13}$C) | 130 mg/L |

#### References


1. C. A. Waudby, A. Ramos, L. D. Cabrita, J. Christodoulou, Two-Dimensional NMR Lineshape Analysis. *Scientific Reports* **6**, 24826 (2016).
2. D. Lee, C. Hilty, G. Wider, K. Wüthrich, Effective rotational correlation times of proteins from NMR relaxation interference. *Journal of Magnetic Resonance* **178**, 72-76 (2006).
3. N. A. Lakomek, J. Ying, A. Bax, Measurement of 15N relaxation rates in perdeuterated proteins by TROSY-based methods. *J Biomol NMR* **53**, 209-221 (2012).
